# Multidimensional feature tuning in category-selective areas of human visual cortex

**DOI:** 10.1101/2025.06.17.659578

**Authors:** Leonard E. van Dyck, Martin N. Hebart, Katharina Dobs

**Author notes:** equal contribution (Martin N. Hebart, Katharina Dobs).

## Abstract

Human high-level visual cortex has been described in two seemingly opposed ways. A categorical view emphasizes discrete category-selective areas, while a dimensional view highlights continuous feature maps spanning across these areas. Can these divergent perspectives on cortical organization be reconciled within a unifying framework? Using data-driven decomposition of fMRI responses in face-, body-, and scene-selective areas, we identified overlapping activity patterns shared across individuals. Each area encoded multiple interpretable dimensions tuned to both finer subcategory features and coarser cross-category distinctions beyond its preferred category, even in the most category-selective voxels. These dimensions formed distinct clusters within category-selective areas but were also sparsely distributed across the broader visual cortex, supporting both locally selective, category-specific, and globally distributed, feature-based coding. Together, these findings suggest multidimensional tuning as a fundamental organizing principle that integrates feature-selective clusters, category-selective areas, and large-scale tuning maps, providing a more comprehensive understanding of category representations in human visual cortex.

## Introduction

Human high-level visual cortex contains category-selective areas that respond preferentially to specific visual inputs, such as faces, bodies, and scenes (Downing et al., 2001; Epstein & Kanwisher, 1998; Kanwisher, 2010; Kanwisher et al., 1997). A widely used approach contrasts predefined stimulus categories, which are hypothesized to require selective processing, to reveal cortical clusters specialized for distinct domains. This strategy supports a categorical view of cortical organization, which has been instrumental in identifying classical category-selective areas (Epstein & Baker, 2019; Kanwisher & Yovel, 2006; Peelen & Downing, 2007), discovering novel category-related responses (Abassi & Papeo, 2024; Bracci et al., 2010; Cortinovis et al., 2025; Henderson et al., 2025; McCandliss et al., 2003), and establishing their behavioral relevance (Cohen et al., 2017; Kanwisher & Barton, 2011; Moro et al., 2008).

At the same time, growing evidence suggests that category-selective areas encode a broad variety of features beyond their preferred categories (Cichy et al., 2012; Haxby et al., 2001). Within these areas, functional subclusters represent both fine-grained, domain-specific features (Bracci et al., 2015; de Haas et al., 2021; Orlov et al., 2010) and domain-general features shared across categories (Arcaro et al., 2009; Çelik et al., 2021; Çukur et al., 2013, 2016), indicating a more complex and overlapping representational organization. In contrast to the categorical view, a dimensional view proposes that occipitotemporal cortex (OTC), including category-selective areas, is organized along continuous feature dimensions that span categorical boundaries (Bao et al., 2020; Bracci & Op de Beeck, 2023; Haxby et al., 2001, 2011). Supporting this view, neural tuning in OTC has been shown to reflect a wide range of feature dimensions, including animacy, real-world size, aspect ratio, behavioral aspects, and semantic content (Abdel-Ghaffar et al., 2024; Almeida et al., 2023; Arcaro & Livingstone, 2024; Bao et al., 2020; Coggan & Tong, 2023; Contier et al., 2024; Huth et al., 2012; Konkle & Caramazza, 2013; Konkle & Oliva, 2012; Watson et al., 2017; Watson & Andrews, 2024). These features are encoded in large-scale maps that embed category-selective areas within a continuous representational landscape.

Whether high-level visual areas are primarily organized by discrete categories or continuous features remains a central and unresolved question (Op de Beeck et al., 2008; Peelen & Downing, 2017; Ritchie et al., 2024; Yargholi & Beeck, 2023). Each perspective provides valuable insights into the functional organization of OTC. The categorical view emphasizes locally selective, domain-specific coding, while the dimensional view highlights globally distributed, feature-based coding. Reconciling these seemingly divergent accounts may be essential for a more comprehensive understanding of how the brain represents complex visual information.

To address this question, we applied a data-driven decomposition of fMRI voxel responses (Khosla et al., 2022) in face-, body-, and scene-selective areas during natural image viewing (Allen et al., 2022), aiming to uncover their underlying representational dimensions. Our approach builds on two theoretical assumptions (Hebart et al., 2020): first, that neural representations are sparse, such that only a limited number of dimensions are required to capture the response to a given stimulus; and second, that these dimensions are continuous and positive, facilitating interpretability by ensuring that each dimension contributes additively to the overall response. To implement these assumptions, we employed non-negative matrix factorization, a method that identifies part-based, sparse, and interpretable components. This approach allowed us to directly test the two competing views: if the categorical view holds, responses in each area should be dominated by a single component aligned with its preferred category; if the dimensional view is correct, responses should span multiple overlapping dimensions reflecting a continuous feature space.

By analyzing both the representational content and cortical topography of the identified dimensions, we found that both categorical and dimensional views capture essential aspects of high-level visual cortex organization. Category-selective areas encode information along multiple dimensions that are consistent across individuals. Dimensions in these areas are dominated by the respective preferred category but are also tuned to additional features, even in the most category-selective voxels. This tuning reveals representations that are richer and more multifaceted than predicted by a strictly categorical account. The topography of these dimensions reflects locally sparse yet globally distributed coding, with distinct clusters within category-selective areas and widely spaced clusters across cortex that together form large-scale maps (Bogdan et al., 2025; Contier et al., 2024; Ritchie et al., 2024; Weiner & Grill-Spector, 2010). This organization integrates functional preferences across multiple spatial scales, from feature-selective clusters to category-selective areas and large-scale tuning gradients. Within this framework, category-selective areas emerge as a special case of sparse tuning along a limited set of dimensions. We propose multidimensional tuning as a unifying organizing principle that reconciles categorical and dimensional views, offering new insight into how the visual system balances specificity and flexibility in representing the visual world.

## Results

### Data-driven voxel decomposition reveals multiple interpretable dimensions in category-selective areas

If category-selective areas encode information along multiple representational dimensions rather than through strict categorical distinctions, then a data-driven decomposition of their activity patterns should reveal distinct but overlapping dimensions. To test this possibility, we analyzed fMRI voxel responses from participants viewing natural images from the Natural Scenes Dataset (NSD; Allen et al., 2022). Our analysis focused on three well-established category-selective areas associated with high-level visual processing: fusiform face area (FFA; Kanwisher et al., 1997), extrastriate body area (EBA; Downing et al., 2001), and parahippocampal place area (PPA; Epstein & Kanwisher, 1998).

To identify the underlying representations within these areas, we applied Bayesian non-negative matrix factorization (BNMF; Khosla et al., 2022; Schmidt et al., 2009), a technique that decomposes voxel responses into latent dimensions and their corresponding spatial maps. BNMF captures multiple selectivities that may coexist within individual voxels. By enforcing non-negativity, it promotes sparse and part-based representations that are more interpretable than those obtained from conventional dimensionality reduction methods. Moreover, because BNMF does not rely on predefined category labels or anatomical boundaries, it allows for a less biased, data-driven exploration of the functional organization within category-selective cortex. To ensure the robustness and generalizability of our results, we implemented a multi-step analysis pipeline (Fig. 1, see Methods). We first determined the optimal number of latent dimensions for each area using bi-cross-validation designed for matrix factorization (Owen & Perry, 2009; Fig. S1). Next, to account for the stochastic nature of BNMF, we performed multiple decompositions per area and clustered the resulting dimensions within each participant, retaining only robust consensus dimensions. To address inter-individual variability, we then matched dimensions across participants by correlating their response profiles to a shared image set, selecting only those dimensions with consistent patterns across individuals. Finally, to test generalizability, we projected voxel responses to held-out images onto the previously learned dimensions and evaluated how well these projections predicted responses to new images.

**Fig. 1.**
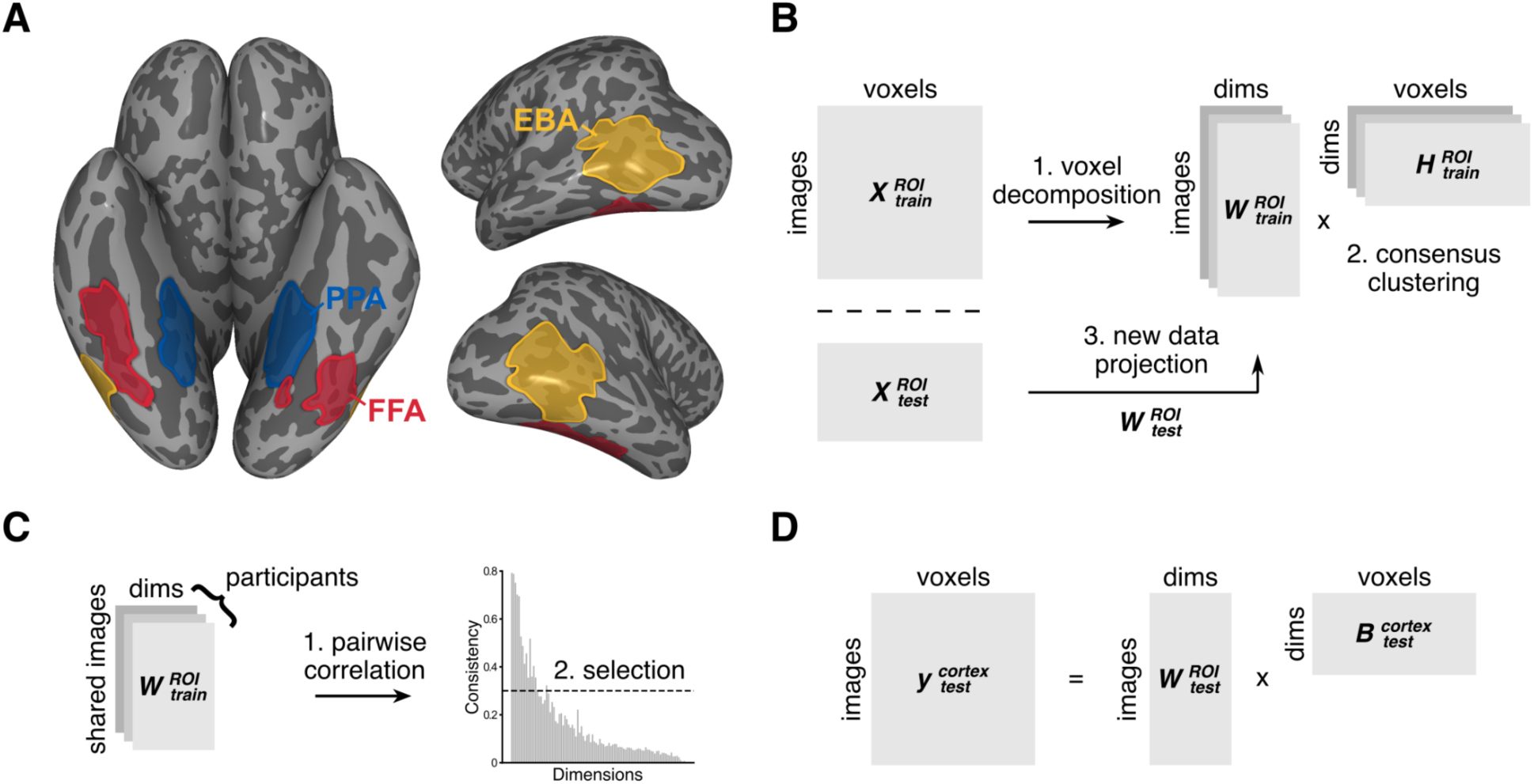
Data-driven fMRI voxel decomposition of category-selective areas. (**A**) FFA, EBA, and PPA were defined using a standard functional localizer (*t* > 2) and are displayed on the inflated cortical surface for a representative participant (P1). (**B**) At the participant level, reliable dimensions were extracted from voxel responses in each area through multiple BNMF decompositions, with the optimal dimensionality determined through bi-cross-validation. Dimensions were clustered across runs using *k*-medoids clustering, and cluster medians were used as reliable consensus dimensions. To assess generalizability, voxel responses to new images were projected onto the learned embedding space using non-negative least squares regression. (**C**) At the group level, consistent dimensions were identified by correlating dimensions across participants for a subset of shared images. Dimensions with the highest inter-participant consistency were retained. (**D**) Voxel-wise encoding models were trained using ridge regression to predict voxel responses to new images based on the learned dimensions, producing functional tuning maps for each dimension across cortex.

Our analysis revealed a small set of highly consistent dimensions in each category-selective area, alongside a larger set of less consistent dimensions, following a function with approximately exponential decay (Fig. 1C, Fig. S2). Despite strong univariate category selectivity (*t* > 2), each area exhibited multiple dimensions that were shared across participants. We identified eight consistent dimensions in FFA, twenty in EBA, and ten in PPA (mean *r* across participant pairs ± *SD*; FFA: *r* = 0.46 ± 0.12; EBA: *r* = 0.49 ± 0.14; PPA: *r* = 0.45 ± 0.10), together capturing a substantial proportion of voxel response variance in each area (mean *R²* across participants ± *SD*; FFA: *R²* = 0.33 ± 0.06; EBA: *R²* = 0.39 ± 0.01; PPA: *R²* = 0.29 ± 0.09). These findings suggest a more nuanced, multidimensional organization than predicted by a strictly categorical account.

### Dimensions reveal multifaceted feature tuning in category-selective areas

Having identified multiple consistent dimensions, we next examined their representational content and how they relate to known functional selectivities. We began by visually inspecting the images that scored highest on each dimension to gain an initial understanding of their semantic content. These images largely aligned with semantic concepts, consistent with the established role of category-selective areas in high-level, view-invariant visual processing. To interpret the dimensions more systematically, we adopted a data-driven labeling approach using multimodal deep learning models. Specifically, we used GPT-4V to generate candidate labels from the highest-scoring images for each dimension and quantified label-image alignment using CLIP-ViT embeddings (Radford et al., 2021). This approach produced meaningful descriptive labels for approximately 95 % of the consistent dimensions (FFA: 8/8; EBA: 20/21; PPA: 9/10; Fig. 2; Fig. S3), although a few remained ambiguous and were left unlabeled.

**Fig. 2.**
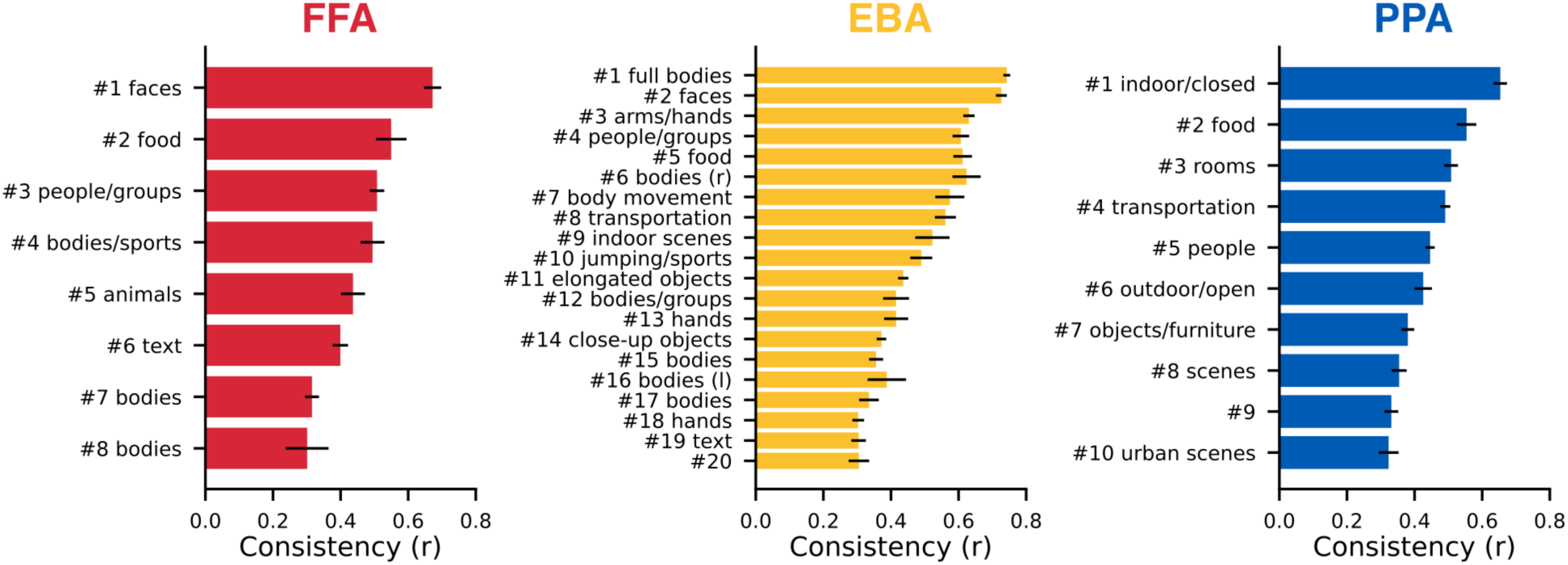
Consistency of dimensions in category-selective areas. Mean correlation of dimensions in each area, averaged across participant pairs (± *SEM*). The labels were determined using a data-driven interpretation based on multimodal deep learning models.

As expected, the most consistent dimensions in each area aligned with its preferred category: faces in FFA, whole bodies in EBA, and scenes in PPA (Fig. 3). However, the dimensions were not limited to broad categories but also captured finer-grained subcategory features. In EBA, for example, dimensions differentiated body parts, including full bodies, arms, hands, and faces (Bracci et al., 2010, 2015; Downing & Peelen, 2011; Orlov et al., 2010; Ramirez et al., 2024). Similarly, in PPA, dimensions encoded scene properties such as openness and naturalness, distinguishing indoor from outdoor, enclosed from open, and urban from natural environments (Kravitz et al., 2011; Lescroart & Gallant, 2019; Park et al., 2011). In contrast, in FFA, dimensions did not show evidence of subcategory tuning to specific facial parts.

**Fig. 3.**
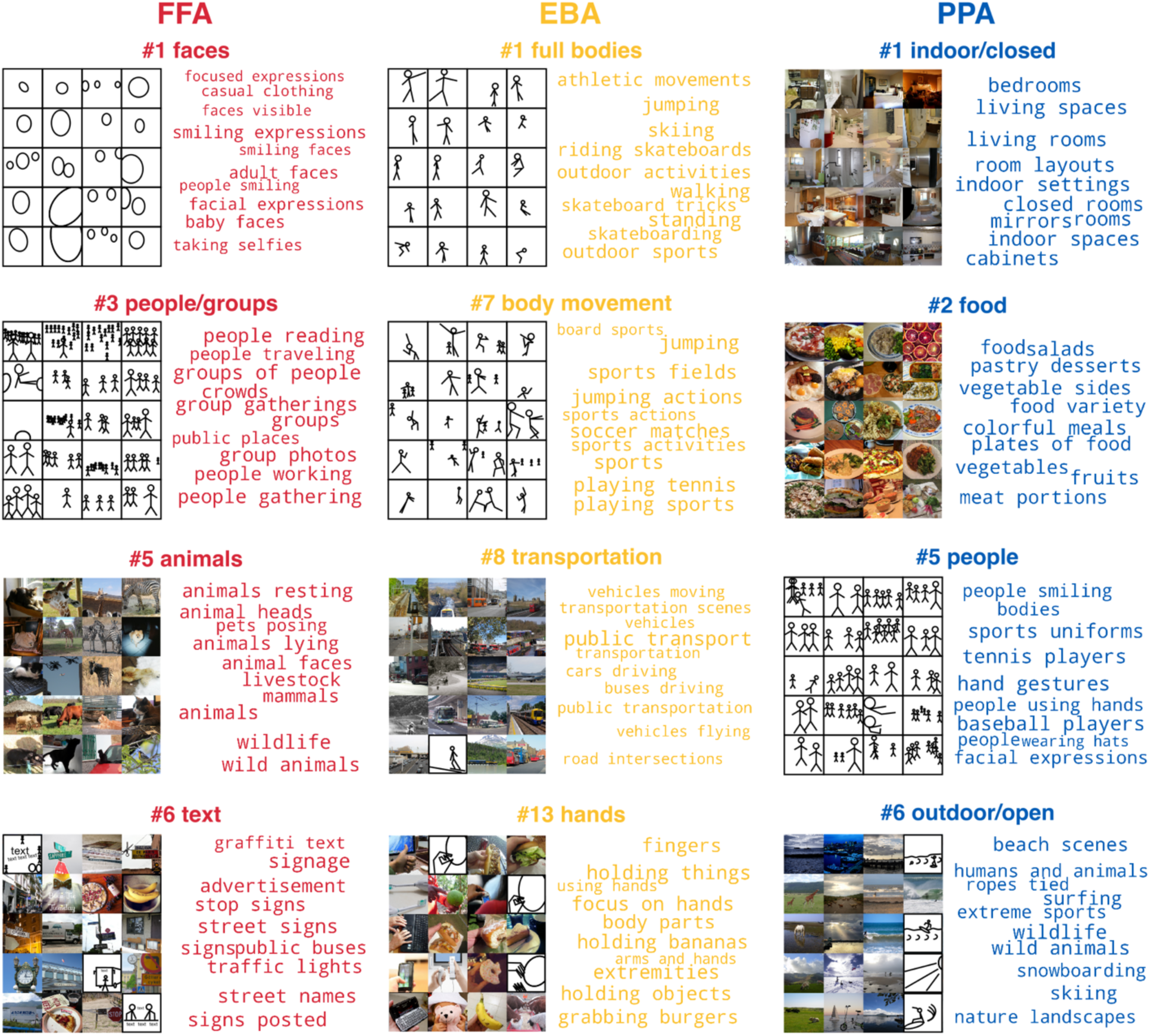
Interpretability of dimensions in category-selective areas. Interpretations of example dimensions from each area. Five highest-scoring images per participant based on dimension response profiles and ten highest-scoring labels combined across participants based on multimodal deep learning models. Images of people were replaced with stick figures.

Beyond dimensions aligned with each area’s preferred category, many additional dimensions captured semantically meaningful content from other domains. In FFA, while some dimensions reflected additional people-related concepts, such as groups, bodies, and sports, others extended to animals, food, and text. In EBA, dimensions were not only tuned to additional body-related concepts, such as movement and sports, but also to food, transportation, indoor scenes, close-up objects, and text. In PPA, alongside additional scene-related concepts, such as transportation and objects/furniture, dimensions also represented food and people. These findings suggest that category-selective areas are not narrowly tuned to a single domain but instead encode a broader semantic space. The lower consistency of these additional dimensions, relative to the dominant category-aligned ones, may help explain why previous approaches were unable to detect this more nuanced and multifaceted tuning in these areas.

Most of the identified dimensions reflected high-level semantic content. A partial exception was observed in EBA, where certain body-related dimensions were tuned to the spatial location of bodies on the left or right side of the image, potentially reflecting retinotopic organization. Apart from this, the dimensions exhibited little sensitivity to low-or mid-level visual features such as shape, texture, or color.

These findings suggest that although category-selective areas show strong tuning to their preferred category at the univariate level, they also encode a broader set of rich, semantic dimensions that become apparent at the multivariate level.

It is possible that identified dimensions within category-selective areas might lie along a single category-selective continuum (e.g., a “faceness” continuum in FFA with faces > bodies > text) rather than representing distinct tuning axes (e.g., faces, bodies, and text in FFA). To address this concern, we analyzed pairwise cosine similarities between dimension response profiles within each area. We specifically compared two groups of dimensions: those tuned to each area’s preferred category, and those tuned to non-preferred features. A single category-selective continuum would predict high similarities between “preferred” and “non-preferred” dimensions (between-group comparison), due to alignment with the dominant category-selective axis. However, an area may also respond preferentially to some stimuli over others, irrespective of category preference, which would be indicated by high similarities among non-preferred dimensions (within-group comparison). Using raw dimension response profiles, we observed moderately high between-group similarities (mean cosine similarity across participants *± SD*; FFA: *M* = 0.66 *±* 0.05; EBA: *M* = 0.64 *±* 0.03; PPA: *M* = 0.72 ± 0.02; range: [0, 1]), as well as moderately high within-group similarities for both preferred (FFA: *M* = 0.65 *±* 0.06; EBA: *M* = 0.72 *±* 0.02; PPA: *M* = 0.78 ± 0.03) and, critically, also non-preferred dimensions (FFA: *M* = 0.67 ± 0.05; EBA: *M* = 0.65 ± 0.03; PPA: *M* = 0.71 ± 0.02). This demonstrates that the generally high similarity observed between dimensions within an area cannot be explained by category-selective preferences. Next, to test if there is residual evidence of category-selective continuum beyond the overall preference for certain stimuli over others, we controlled for the overall response increase by removing the mean response profile across images and recomputing the similarities (Fig. S4). As expected, after mean-centering, we observed low between-group similarities (FFA: *M* =-0.13 *±* 0.02; EBA: *M* =-0.11 *±* 0.01; PPA: *M* =-0.14 ± 0.01; range: [-1, 1]), as well as low within-group similarities for both preferred (FFA: *M* =-0.14 ± 0.05; EBA: *M* = 0.01 ± 0.01; PPA: *M* =-0.02 ± 0.03) and non-preferred dimensions (FFA: *M* =-0.13 ± 0.01; EBA: *M* = 0.02 ± 0.01; PPA: *M* =-0.10 ± 0.01), in line with multidimensional tuning. These results show that the dimensions not only show common stimulus preferences, but also complementary (between-group) and distinct (within-group) tuning patterns that encode multiple, partially independent tuning axes rather than a single category-selective continuum.

Together, these findings demonstrate that category-selective areas encode rich, multifaceted representations that extend beyond simple categorical boundaries. These areas capture both finer subcategory features and broader cross-category distinctions, underscoring the complexity and flexibility of functional organization in high-level visual cortex.

### Multidimensional tuning underlies category-selective representations

A potential concern is that the observed multidimensional tuning might simply arise from the inclusion of weakly selective voxels in the definition of category-selective areas. If so, the apparent diversity in tuning could reflect a methodological artifact rather than a genuine feature of functional organization. To address this possibility, we examined whether a voxel’s degree of category selectivity predicted its tuning specificity across dimensions. For each voxel, we quantified category selectivity using its preferred category contrast from the functional localizer (d’; e.g., faces > all other categories in FFA) and related this measure to the voxel’s weights across the learned dimensions. We also computed a sparseness measure (Hoyer, 2004; see Methods) to quantify each voxel’s tuning profile, with high sparseness indicating narrow tuning to only a few dimensions and low sparseness reflecting broader tuning to multiple dimensions. Across all areas, we found no evidence that greater category selectivity was associated with narrower dimensional tuning. Voxel-wise category selectivity was not reliably related to sparseness in EBA or PPA and showed only a trend in FFA (mean *r* across participants ± *SD*; FFA: *r* = 0.23 ± 0.15, *p* = 0.06; EBA: *r* =-0.03 ± 0.07, *p* = 0.46; PPA: *r* =-0.01 ± 0.08, *p* = 0.77; Fig. S5). These results suggest that even the most category-selective voxels remain tuned to multiple, partially overlapping dimensions.

Further analysis revealed that although dominant dimensions in highly category-selective voxels corresponded primarily to the preferred category, additional dimensions captured complementary features (Fig. 4). For example, highly face-selective voxels in FFA also responded to dimensions related to bodies and animals, highly body-selective voxels in EBA were additionally tuned to actions, faces, and elongated objects, and highly scene-selective voxels in PPA responded to dimensions related to transportation, objects/furniture, and people.

**Fig. 4.**
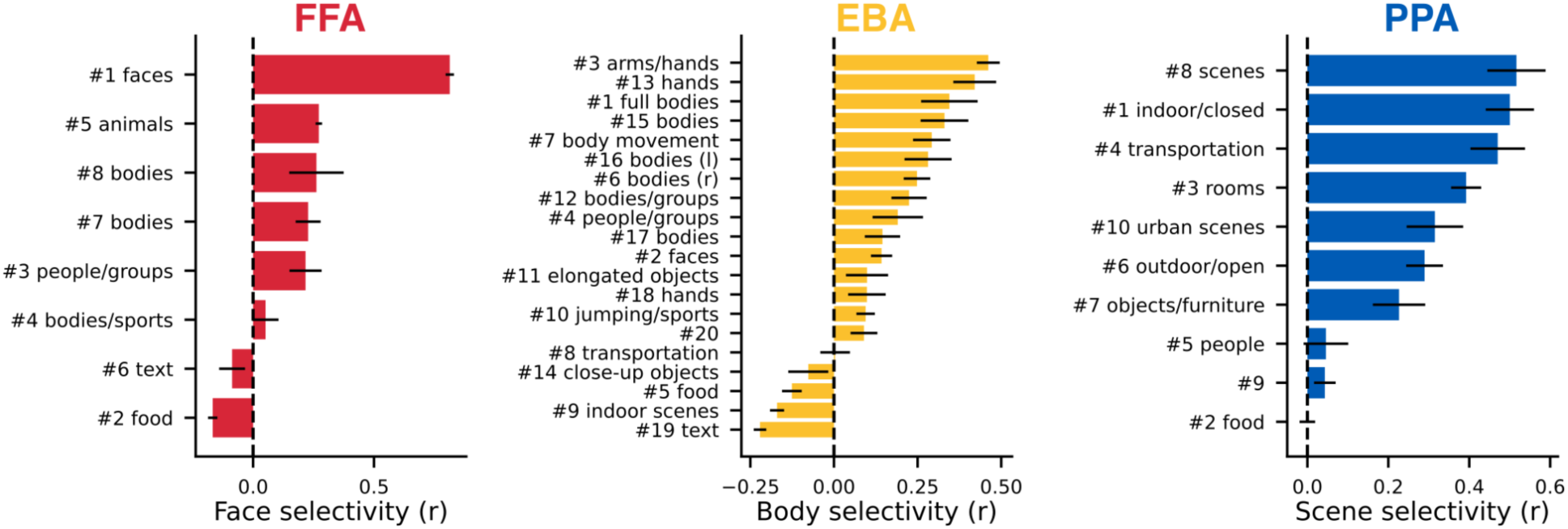
Relationship between category selectivity and dimension tuning. Mean correlation of voxel-wise dimension weights and category selectivity based on the respective functional localizer contrast in each area, averaged across participants (± *SEM*).

Together, these findings indicate that multidimensional tuning is not merely a byproduct of weak voxel selectivity. Even the most category-selective voxels exhibit tuning to multiple dimensions. This suggests that category selectivity does not result from exclusive tuning to a single feature axis, but rather from a multidimensional code that integrates information across semantically related feature axes.

### Dimensions from category-selective areas explain activity throughout high-level visual cortex

Having characterized the representational content of the identified dimensions, we next investigated whether these dimensions were confined to specific cortical areas or distributed more broadly across cortex. To test this, we assessed how well dimensions derived from each category-selective area predicted voxel responses beyond their region of origin. If the representations were highly localized, predictive power would be restricted to the source area. In contrast, broader topographies would suggest a more distributed functional organization with overlapping and shared dimensions.

We addressed this question using voxel-wise encoding models trained to predict responses to natural images based on the dimensions derived from each category-selective area. Model performance was evaluated by predicting voxel responses to held-out test images across cortex. To accommodate the possibility that some dimensions negatively correlate with neural activity elsewhere, we relaxed the non-negativity constraint on the voxel-wise coefficients. This allowed us to map the prediction performance of each dimension set and evaluate their explanatory power across the cortical surface (Fig. 5, Fig. S6). Prediction performance was highest in high-level visual areas, consistent with the semantic nature of the dimensions, but also extended into early visual areas and prefrontal cortex, as confirmed through permutation testing. Within each category-selective area, locally derived dimensions explained local tuning especially well, with performance often approaching the estimated noise ceiling (Fig. S7). Importantly, the dimensions generalized across areas. For example, FFA-derived dimensions explained variance in body-selective areas such as EBA and fusiform body area (FBA), while EBA-derived dimensions predicted responses in face-selective areas such as FFA and superior temporal sulcus (STS). Similarly, EBA-derived dimensions predicted responses in scene-selective areas such as PPA and occipital place area (OPA). While some generalization may be attributable to spatial overlap between adjacent areas (e.g., FFA and FBA, EBA and OPA), the strength of the cross-predictive performance suggests that shared representational dimensions, rather than strict categorical boundaries, underlie these relationships.

**Fig. 5.**
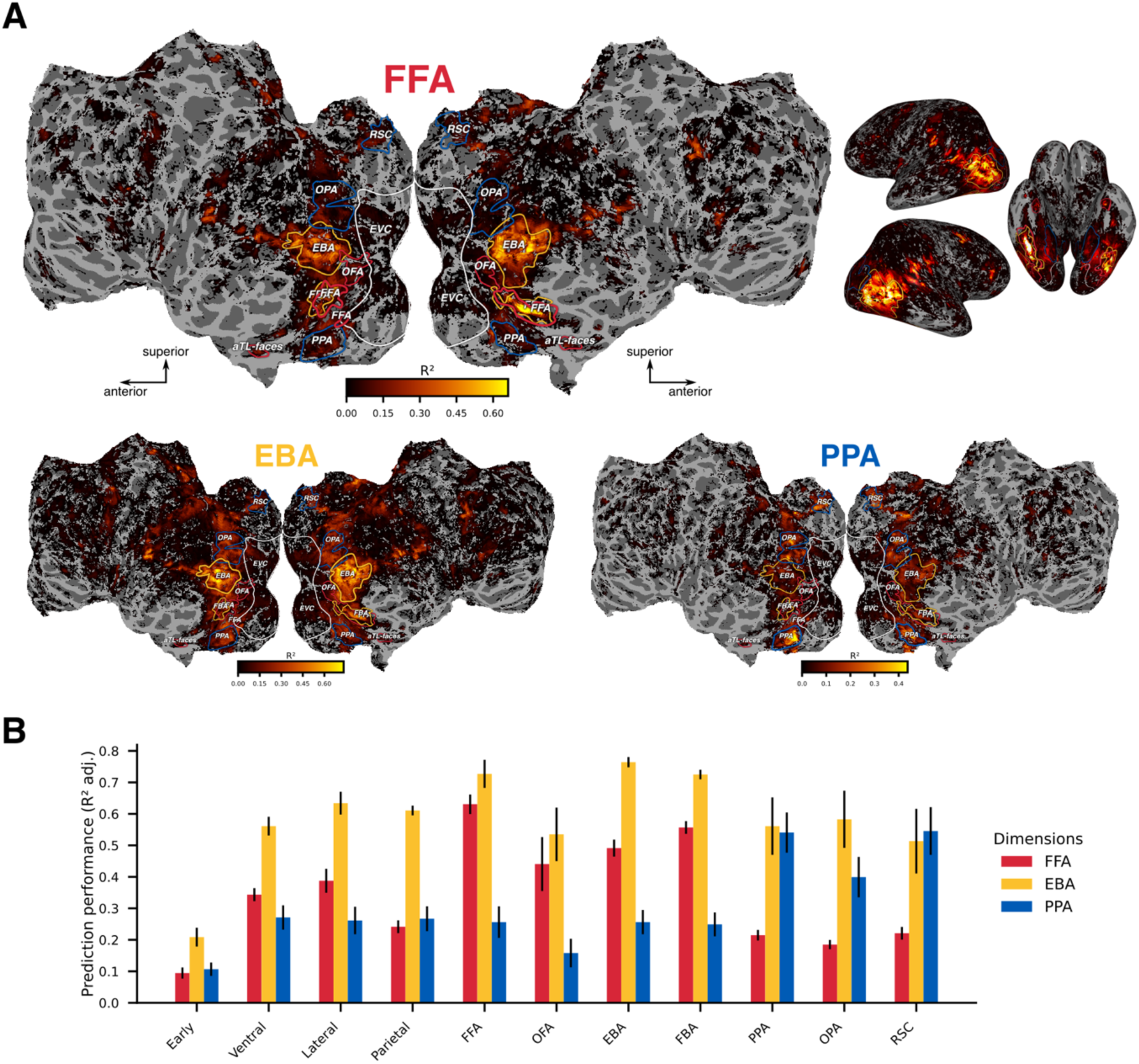
Prediction performance of dimensions from category-selective areas across cortex. (**A**) Voxel-wise prediction performance (*R^2^*) for dimensions from each area projected onto the flattened cortical surface for a representative participant (P1). Voxels thresholded at *p* < 0.01 (one-sided, FDR-corrected). (**B**) Mean prediction performance (noise-ceiling-adjusted *R^2^*) for different cortical areas, averaged across participants (± *SEM*).

Together, these findings indicate that while the dimensions capture the specific tuning profiles of individual category-selective areas, they also reveal a more distributed and overlapping functional organization. Rather than operating as isolated modules, these areas appear to participate in broader cortical networks that encode both unique and shared features.

### Dimensions form locally sparse clusters and globally distributed maps

Having established that dimensions derived from category-selective areas explain responses across broad regions of cortex, we next examined their individual topographies. Within category-selective areas, the spatial organization of these dimensions could reflect either broad overlap, consistent with functional homogeneity, or discrete clusters, consistent with functional heterogeneity. To distinguish between these possibilities, we visualized the voxel-wise coefficient weights for each dimension across the flattened cortical surface (Fig. 6, Figs. S8-11).

**Fig. 6.**
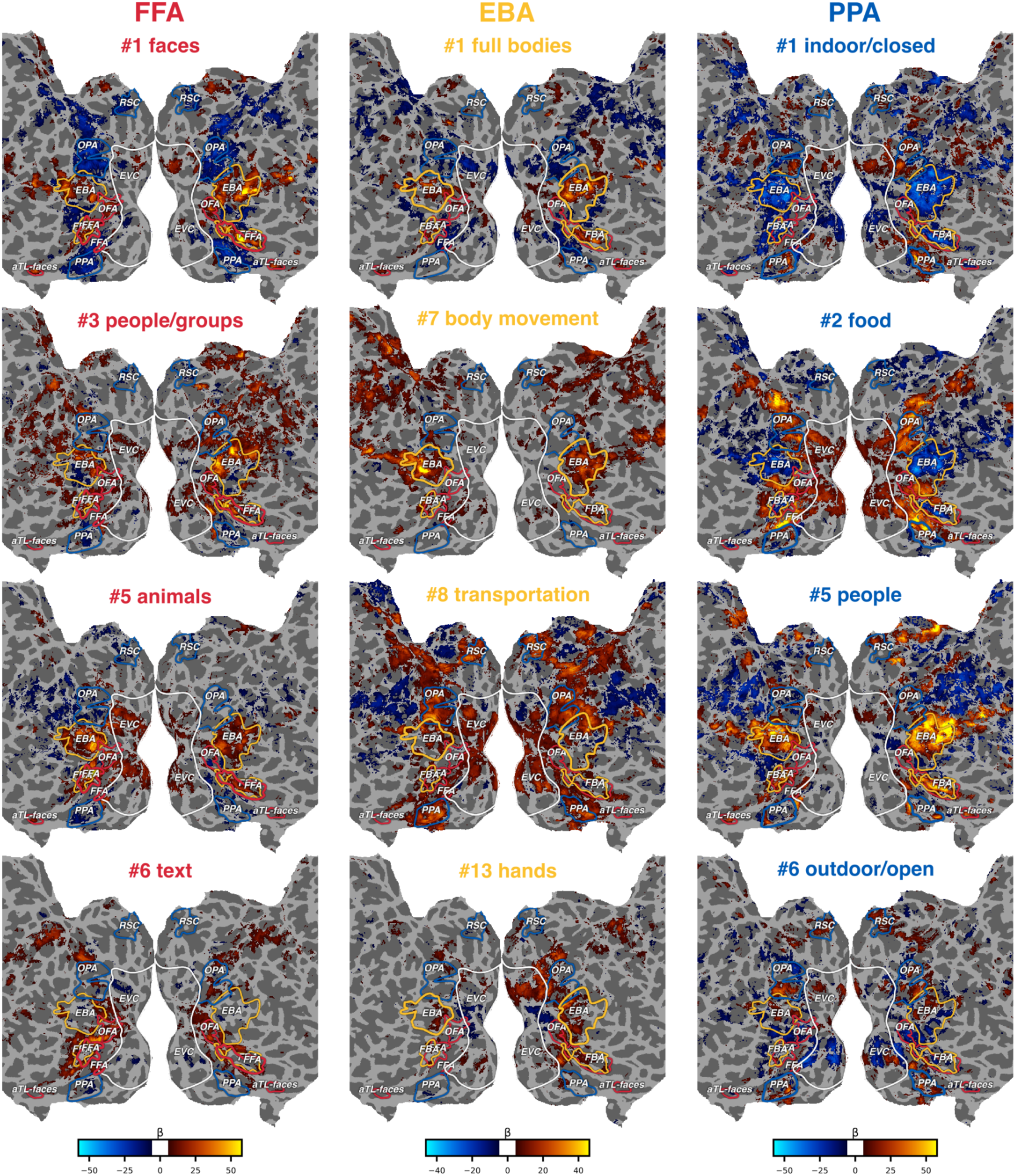
Functional tuning maps of individual dimensions from category-selective areas across cortex. Voxel-wise coefficient weights for six example dimensions from each area projected onto the flattened cortical surface for a representative participant (P1). Voxels thresholded at *p* < 0.01 (one-sided, FDR-corrected).

Many dimensions reflected known functional selectivities: face-related dimensions were concentrated in face-selective areas, body-related dimensions in body-selective areas, and scene-related dimensions in scene-selective areas. However, these representations were not spatially uniform. Most dimensions formed distinct subclusters within each category-selective area, revealing a finer-grained functional organization. In FFA, subclusters were tuned to non-face features (Çukur et al., 2013). In EBA, subclusters encoded different body parts, including full bodies, faces, arms, and hands (Bracci et al., 2010, 2015; Downing & Peelen, 2011; Orlov et al., 2010; Ramirez et al., 2024). In PPA, subclusters encoded different scene properties, such as openness and possibly distance (Lescroart & Gallant, 2019). These findings extend earlier reports that category-selective areas encode non-preferred features (Cichy et al., 2012; Haxby et al., 2001), demonstrating that such representations are not only present but also spatially organized. This supports an account in which category-selective areas are composed of multidimensional tuning maps rather than uniform modules, consistent with prior proposals of finer-grained functional organization (Grill-Spector & Weiner, 2014). Some dimensions also revealed novel or under-characterized selectivities, such as tuning to animals in FFA and elongated objects in EBA. These may reflect finer subdivisions within areas and highlight the potential of this approach to uncover new functional specializations.

Beyond local clustering, many dimensions extended across area boundaries and overlapped with neighboring regions. For example, in FFA, a text-related dimension extended into visual word form area (VWFA), and body-related dimensions overlapped with FBA. Other dimensions, such as those tuned to food, were even more broadly distributed, spanning multiple areas like V4, FFA, and PPA (Contier et al., 2024; Jain et al., 2023; Khosla et al., 2022; Pennock et al., 2023). We also observed expected hemispheric asymmetries, with face-and body-related dimensions predominantly right-lateralized and text-related dimensions left-lateralized (Dehaene & Cohen, 2011; Willems et al., 2010). Overall, the cortical organization of these dimensions was marked by locally sparse clustering and globally distributed tuning across high-level visual cortex (Bogdan et al., 2025; Contier et al., 2024; Miyakawa et al., 2018; Ritchie et al., 2024; Weiner & Grill-Spector, 2010). This organization revealed functionally selective clusters distributed across cortex. For example, an EBA-derived face dimension recurred in other face-selective areas, including FFA, STS, occipital face area (OFA), an anterior temporal face area (aTL-faces), and a prefrontal face area near inferior frontal sulcus (Nikel et al., 2022; Tsao et al., 2008). Similarly, people-related dimensions were mostly shared between EBA and FFA, supporting the existence of interconnected face-and body-selective networks (Taubert et al., 2022; Weiner & Grill-Spector, 2010). In addition, body-related dimensions were found in and around multiple scene-selective areas, as recently reported regarding the existence of new body-selective areas (Zhao et al., 2025).

To determine whether these overlapping functional maps also capture large-scale tuning gradients, we analyzed the major tuning directions using principal component analysis of the dimension profiles. This approach provided a coarse summary of the dominant tuning patterns and their corresponding spatial gradients, revealing that multidimensional tuning mirrors previously reported large-scale gradients (Çelik et al., 2021; Huth et al., 2012; Konkle & Caramazza, 2013; Fig. S12). Despite some inter-individual variability in spatial location, the overall arrangement was consistent across participants, suggesting that these dimensions reflect stable and generalizable principles of cortical organization.

In summary, our findings provide a comprehensive account of functional organization in high-level visual cortex. Individual dimensions form sparse subclusters within category-selective areas while also contributing to distributed, large-scale maps across cortex. This locally sparse but globally distributed organization supports both discrete category selectivity and continuous feature integration, offering a unified framework for how high-level visual cortex encodes complex information.

## Discussion

Is high-level visual cortex organized into discrete categorical modules or does it reflect a continuous representational space? Our data-driven analysis of fMRI responses suggests it is neither strictly one nor the other, but a hybrid: a multidimensional code that integrates elements of both discrete and continuous organization. Within category-selective areas, we identified interpretable dimensions that largely aligned with their preferred categories, consistent with domain-specific tuning (Downing et al., 2005; Kanwisher, 2010). At the same time, these areas also encoded finer subcategory features and coarser cross-category distinctions, uncovering more nuanced and multifaceted representational content. These findings are consistent with recent evidence that category-selective areas flexibly encode both domain-specific and domain-general information (Khosla & Wehbe, 2023; Vinken et al., 2023).

Our study contributes to the understanding of high-level visual cortex organization in several ways. Despite being based directly on cortical responses to complex natural images, the decomposition yielded remarkably interpretable dimensions. Combining this decomposition with encoding models allowed us to link functional hallmarks across multiple spatial scales, ranging from fine-scale subclusters, over category-selective areas, to large-scale cortical maps. Moreover, our results showed that even the most category-selective parts of high-level visual cortex are tuned to multiple meaningful dimensions, calling into question the notion of pure selectivity at the voxel level.

We also found that representational diversity differed across category-selective areas. While FFA exhibited the narrowest tuning, both EBA and PPA displayed broader and more heterogeneous tuning. Notably, EBA showed the greatest heterogeneity, consistent with its role in encoding individual body parts (Bracci et al., 2015; Downing & Peelen, 2011; Orlov et al., 2010; Ramirez et al., 2024) and its spatial overlap with multiple anatomically and functionally defined subregions (Weiner & Grill-Spector, 2011). These findings suggest that EBA may play a more integrative role than traditionally assumed. Similarly, PPA encoded scene-related features such as naturalness, openness, and presumably distance (Kravitz et al., 2011; Lescroart & Gallant, 2019; Park et al., 2011). In contrast, FFA did not show separable tuning to distinct facial parts, as observed in more controlled studies (de Haas et al., 2021; Henriksson et al., 2015), possibly due to limited variability or smaller size in the natural images. Interestingly, we were able to identify selectivity to animals in FFA, which may reflect sensitivity to animal faces or broader animacy features (Kanwisher et al., 1999).

In addition to identifying individual dimensions, we uncovered two key principles of their spatial organization. First, within category-selective areas, dimensions formed distinct subclusters, consistent with a mosaic-like organization of feature-selective populations (Çukur et al., 2013, 2016; Grill-Spector & Weiner, 2014). Second, many dimensions were sparsely distributed across high-level visual cortex, suggesting a general-purpose representational scaffold. Together, these findings support a hierarchical organization, where fine-scale subclusters are tuned to specific dimensions, category-selective areas are sparsely tuned to subsets of these dimensions, and large-scale cortical maps reflect integrative, overlapping codes.

Our findings resonate with a growing body of work using encoding models to uncover nuanced patterns of functional organization in the brain (Çelik et al., 2021; Doerig et al., 2022; Efird et al., 2024; Luo et al., 2024). Notably, the topographies of individual dimensions closely resembled those of previously reported behavior-derived object dimensions (Contier et al., 2024; Hebart et al., 2020), suggesting that these neural representations may contribute to goal-directed behavior.

Building on this work, our results support the idea that category selectivity emerges from sparse tuning to a limited number of high-level dimensions (Contier et al., 2024; Ritchie et al., 2024). This view is consistent with prior evidence linking category selectivity to sparsely distributed neural codes and connectivity-based constraints (Miyakawa et al., 2018; Molloy et al., 2024; Op de Beeck et al., 2019; Osher et al., 2016; Weiner & Grill-Spector, 2010). In contrast, other accounts propose that category selectivity arises from tuning to many directions within a dense, high-dimensional feature space shaped by mid-level visual properties (Vinken et al., 2023), making it a possible outcome of broadly distributed coding rather than a special case of sparse tuning. Recent computational work offers a possible route for the emergence of multidimensional tuning, showing that contrastive learning can produce brain-like functional organization in neural network models without relying on explicit category-specific mechanisms (Prince et al., 2024). Similarly, topographic neural network models offer a useful framework for investigating how overlapping feature tuning might produce spatial patterns that integrate selectivity across scales (Blauch et al., 2022, 2025; Doshi & Konkle, 2023; Lu et al., 2025; Margalit et al., 2024). Additionally, emerging evidence suggests that such dimensions may also be dynamically modulated over time (Shi et al., 2023; Teichmann et al., 2024). Together, these findings support an integrative account in which category selectivity arises from spatially organized tuning and possibly temporally dynamic modulation of a small number of high-level dimensions embedded in a broader representational space.

Despite its contributions, our approach has several limitations. First, our decomposition approach imposes a non-negativity constraint, which is well-suited for capturing excitatory BOLD responses but insensitive to potentially meaningful inhibitory signals (Pérez-Ortega et al., 2024). Although we relaxed this constraint in the encoding models, future work should explore more flexible techniques that capture both excitatory and inhibitory components. Second, while many of the identified dimensions were interpretable, some remained ambiguous, likely reflecting many-to-one mappings between visual features and neural activity. Accordingly, the descriptive labels generated via our data-driven deep learning approach should be viewed as hypotheses rather than definitive interpretations. Third, like any data-driven approach, our findings are shaped by the properties of the dataset. Limited diversity in the natural images may bias the recovered dimensions (Shirakawa et al., 2025), although our focus on robustness likely mitigated the influence of spurious effects. Nonetheless, disentangling stimulus-driven features from learned associations remains a central challenge when using complex natural stimuli.

Our findings point to several promising directions for future research. A central question is whether these dimensions guide behavior and how they adapt to varying task demands (Bracci & Op de Beeck, 2023; Peelen & Downing, 2017; Ritchie et al., 2024). Examining their stability across tasks, datasets, and imaging modalities will provide deeper insights into their generalizability. Future work should also investigate how these dimensions interact across areas (Op de Beeck et al., 2008) and evolve over time (Shi et al., 2023; Teichmann et al., 2024), potentially revealing their computational roles. In addition, probing the granularity of these dimensions may clarify how fine-and coarse-scale representations coexist (Gauthaman et al., 2024; Han & Bonner, 2025). While our decomposition showed mixed selectivity at the voxel level, discrete tuning may exist at finer scales, highlighting the need for higher-resolution methods to uncover potentially hidden structure (Quiroga et al., 2005). Finally, comparing biologically derived dimensions to those learned by artificial neural networks may uncover shared representational strategies between biological and artificial vision systems (Chen & Bonner, 2024; Hosseini et al., 2024; Huh et al., 2024; Kanwisher et al., 2023; Mahner et al., 2025).

To conclude, our multidimensional framework bridges the divide between categorical and dimensional accounts of high-level visual cortex organization. Our findings demonstrate that, when analyzed in a data-driven manner, these perspectives are not mutually exclusive but rather complementary. Category-selective areas emerge from overlapping, sparsely distributed representational dimensions. This framework captures both the apparent discreteness of category selectivity and the continuity of dimensional coding. Multidimensional tuning provides a unifying principle that enables the visual system to balance specificity with flexibility, which is essential for supporting a wide range of functional demands.

## Methods

### fMRI data

#### Natural Scenes Dataset

We used the Natural Scenes Dataset (NSD; Allen et al., 2022), which includes fMRI responses from eight participants who each viewed 9,000 to 10,000 natural scene images over 30 to 40 scan sessions, with each image repeated three times. Our analyses focused on participants who completed all trials (P1: S1, P2: S2, P3: S5, and P4: S7), viewing 9,000 unique and 1,000 shared images. During scanning, participants maintained central fixation and performed a long-term recognition task by identifying previously viewed images. Images were sourced from the Microsoft Common Objects in Context (COCO) database (Lin et al., 2014) and displayed at a visual angle of 8.4° × 8.4° for 3 s, with a 1 s interval between images. BOLD responses were acquired at 7 T using whole-brain gradient-echo EPI with 1.8 mm isotropic voxel resolution and a 1.6 s repetition time. Preprocessing included temporal and spatial interpolation for slice timing and head motion correction, and single-trial voxel responses were estimated using a general linear model. We used preprocessed single-trial voxel responses optimized through voxel-wise hemodynamic response function modeling, data-driven denoising, and ridge regression (“betas_fithrf_GLMdenoise_RR”; 1.8 mm isotropic native preparation; Prince et al., 2022). Cortical flat maps were generated using PyCortex (Gao et al., 2015). Details on scanning protocols, preprocessing steps, and noise ceiling estimation are provided in the original NSD paper (Allen et al., 2022). To cross-validate our analyses, we randomly divided the images into a 70 % training set and a 30 % test set, ensuring that all shared images were assigned to the training set. To account for session effects and improve reliability, single-trial voxel responses were *z*-scored within each session using the mean and standard deviation of the training trials, and neural response estimates were averaged across repetitions of each image.

#### Category-selective areas

We extracted voxel responses from three functional regions of interest (ROIs): fusiform face area (FFA), parahippocampal place area (PPA), and extrastriate body area (EBA). These ROIs were identified using a separate functional localizer (fLOC; Stigliani et al., 2015). During the fLOC, participants viewed grayscale images of five categories (faces, bodies, places, characters, objects) presented in a miniblock design across six runs, with each block containing eight images per category. After standard preprocessing, category-specific beta values were estimated using a general linear model. ROIs were defined by conducting *t*-tests comparing each category to all others (*t* > 2), isolating voxels with clear category preferences. Details on the analyses are provided in the original fLOC paper (Stigliani et al., 2015). To ensure reliable input for BNMF, we selected voxels within each ROI that had a signal-to-noise-ratio greater than 0.2. We also performed the BNMF procedure for additional face-, body-, and scene-selective ROIs, including occipital face area (OFA), fusiform body area (FBA), and occipitotemporal place area (OPA). These analyses yielded results similar to those observed in the primary ROIs (FFA, EBA, and PPA). For the sake of brevity, we focus on the three best-known ROIs here. We quantified voxel-wise category selectivity using the category selectivity index (*d’,* Eq. 1), computed based on each ROI’s preferred category contrast (i.e., preferred category versus all other categories), where 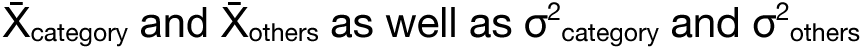 denote the mean voxel responses and variances to the preferred category and all other categories, respectively:

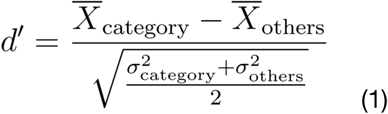

### Data-driven voxel decomposition

#### Bayesian non-negative matrix factorization

We applied Bayesian non-negative matrix factorization (BNMF) to decompose voxel responses from ROIs into dimensions, following prior work by Khosla et al. (2022). Non-negative matrix factorization (NMF) decomposes a data matrix *V* (*I* × *V*), where *I* is the number of images and *V* is the number of voxels, into the product of two lower-rank matrices: a response matrix *W* (*I* × *D*) and a weight matrix *H* (*D* × *V*), where *D* is the number of dimensions, such that the product of these matrices optimally reconstructs the data. In this formulation, each column of *W* (*w_d_*) represents a dimension’s response profile, while each row of *H* (*h_d_*) encodes its spatial pattern across voxels. The non-negativity constraint in NMF restricts dimensions to additive combinations, promoting part-based, sparse, and interpretable dimensions. Compared to other dimensionality reduction techniques like principal component analysis or cluster analysis, NMF offers several advantages: (1) voxel responses are represented exclusively by additive dimensions, avoiding cancellation effects, (2) voxels are not assigned only to individual dimensions or individual clusters, allowing for mixed selectivity, and (3) dimensions are sparse, with each voxel associated with only a subset of dimensions.

We applied BNMF, an extension of NMF, which incorporates Bayesian inference (Eq. 2) by imposing exponential priors on *W* (Eq. 3) and *H* (Eq. 4), and a normal likelihood for the residual matrix *E* (*I* × *V*) with noise variance *σ^2^*. BNMF employs Gibbs sampling to iteratively sample from the posterior distributions of *W*, *H*, and *σ^2^*. This Bayesian formulation yields probabilistic estimates that explicitly model noise in the data, improving robustness. Specifically, BNMF is based on the following model components:

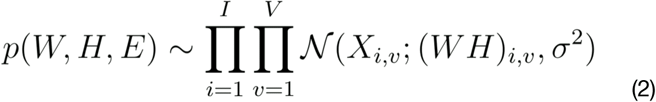

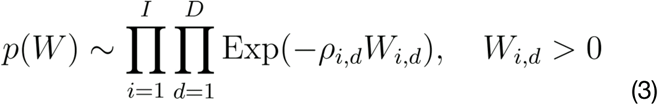

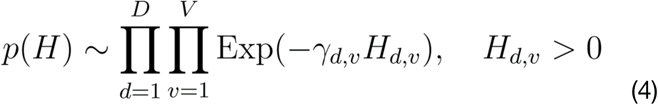

We ran 3,000 Gibbs iterations, discarded the first 1,000 as burn-in, and retained every 5th draw after this. The retained samples were averaged element-wise to obtain the posterior mean estimates of the factor matrices. Each model was initialized with a unique random seed, and convergence was achieved once the reconstruction error fell below 1×10⁻⁵. Details of the algorithm are provided in the original BNMF paper (Schmidt et al., 2009). To enforce non-negativity, we baseline-shifted *X* by subtracting the minimum *z*-scored response across all voxels in each ROI. To prevent data leakage, both training and test sets were shifted using the global training minimum. Any negative outliers remaining in the test set were set to 0. The constant offset introduced between training and test distributions was accounted for in the final encoding model by fitting an intercept term.

#### Estimating the optimal dimensionality

We determined the optimal dimensionality *k\** for each participant’s ROI using bi-cross-validation, which accounts for the interdependence of rows and columns in matrix factorization (Owen & Perry, 2009). The data matrix *X* was shuffled and divided into four blocks: *A*, *B*, *C*, and *D*. BNMF was trained on the training block *D*, and pseudo-inverse matrices of *W* and *H* from blocks *B* and *C* were used to predict the held-out test block *A*. This process was repeated 5 times with random block partitions in each iteration for ranks between 1 and 100 in steps of 3, and the average prediction error across iterations was used to calculate the final cross-validation error. For each participant, *k\** was selected as the rank that minimized the test error (Fig. S1). Estimating the optimal dimensionality for fMRI responses to natural images is challenging due to their hierarchical nature, which requires capturing meaningful dimensions while avoiding overfitting to spurious ones. Our approach balanced these considerations, ensuring optimal latent dimensionality within each participant’s ROI, maximizing representational capacity, while also minimizing the risk of overfitting.

#### Consensus approach

We identified dimensions within each ROI using a two-step approach to ensure reliability and generalizability. First, to address the sensitivity of BNMF to initialization, we adapted a consensus approach outlined in previous work (Kotliar et al., 2019). For each ROI, we performed 100 randomly initialized BNMF voxel decompositions with *k\** dimensions. Each dimension was normalized to an *L_2_*-norm of 1. Unreliable runs were removed based on a density-based outlier detection approach. To aggregate reliable dimensions across runs, we applied *k*-medoids clustering with cosine distance, using *k*\* clusters. Cluster medians were then used as consensus dimensions for subsequent analyses. Second, to identify consistent dimensions across participants, we used a pairwise correlation approach also outlined in previous work (Khosla et al., 2022). We computed pairwise correlations between all combinations of dimensions across participants (*k*_1_* × *k*_2_* × *k*_3_* × *k*_4_*). Using a greedy selection approach, we iteratively matched dimensions with the highest pairwise correlations until no more dimensions could be matched across all participants. Dimensions with a mean pairwise correlation exceeding *r* greater than 0.3 were considered consistent, resulting in 8 to 20 consensus dimensions per ROI (Fig. S2). This two-step approach ensured that the identified dimensions were reliable across runs and consistent across participants, providing a robust framework for subsequent analyses. To evaluate the generalizability of the dimensions, we projected the test set into the learned embedding using non-negative least squares regression.

### Data-driven dimension labeling

To interpret the consistent dimensions identified across participants, we employed a data-driven labeling approach using large language models to generate labels and multimodal neural network representations to align image-related dimensions with meaningful semantic labels. The approach involved generating a diverse set of candidate labels based on the highest-scoring images for each dimension and evaluating the labels using all training images. First, candidate labels were generated using GPT-4V (OpenAI). For each dimension, we selected the five highest-scoring images from each participant and provided these images combined as input to the model, along with the instruction to generate ten concise labels (with a maximum of three words each) per set of images (see Supplementary Text). We pooled the labels generated by GPT-4V across dimensions to compile a comprehensive set of candidate labels. Next, the candidate labels were encoded using CLIP-ViT-L-14 (Radford et al., 2021), with embeddings generated for each label using 15 different prompt templates (e.g., “a picture of…”, “an image of…”, “a photo of…”), following the advised methodology to enhance semantic robustness. Similarly, image embeddings were extracted for all training images. To assess the alignment of each label with the images, we calculated the cosine similarity between the label and image embeddings, weighted by the response profile of each dimension. This weighting ensured that the most relevant images for each dimension were prioritized in the analysis. For each label, we averaged the cosine similarities across the different prompts and performed the following steps to ensure specificity and reduce the influence of generic labels. We first subtracted the global mean similarity for each label across all dimensions. This step reduced global biases, allowing similarities to better reflect the distinctive features associated with each label and dimension. After mean subtraction, the weighted similarities were *z*-scored across all labels. *Z*-scoring helped to emphasize labels that were uniquely associated with each dimension, minimizing the impact of labels with consistently high similarities across all dimensions. The *z*-scored similarities were then used to rank the labels for each dimension, with the top-ranked labels visualized as word clouds to provide a concise, interpretable representation of the dimensions. This method allowed us to derive meaningful, semantically relevant labels for each dimension, based on the recurring patterns in the highest-scoring images, while validating them across the entire image set. By using this approach, we ensured that the interpretations of the dimensions were both robust and representative.

### Voxel-wise encoding model

To investigate prediction performance and functional tuning maps of dimensions from each ROI, we implemented a voxel-wise encoding model using its dimensions to predict voxel responses across cortex. Importantly, we used the consistent BNMF dimensions derived from baseline-shifted ROI voxel responses to predict unshifted voxel responses to new images across cortex. To mitigate overfitting and address multicollinearity between dimensions, we applied fractional ridge regression (Rokem & Kay, 2020), which allows for flexible regularization while preserving the sparse and interpretable nature of the dimensions. This regularization technique balances regularized and unregularized coefficient norms via an *α*-fraction. A fraction of 1 corresponds to no regularization (ordinary least squares solution), while a fraction of 0 represents maximum regularization (shrinking all coefficients to zero). The fraction for each voxel was optimized using 10-fold cross-validation of the training set. For each fold, fractions were tuned on 90 % of the training data and validated on the remaining 10 %, sampled from 0.1 to 0.9 in increments of 0.1, and from 0.9 to 1 in increments of 0.01 for higher precision in the less regularized range. The model included z-scoring of predictors within cross-validation folds and an intercept to account for the baseline-shift. We calculated prediction performance (*R²*) for each fold and selected the fraction that yielded the highest average performance per voxel. Voxels in higher-level visual cortex surrounding the ROIs generally required less regularization and achieved higher prediction performance. The final model, trained using optimal fractions, was evaluated on the held-out test set by capturing the relationship between predicted and observed voxel responses. Statistical significance was assessed by generating a null distribution of correlations through 3,000 random permutations of the test set within each fold. Voxel-wise *p*-values were calculated with a one-tailed comparison to the null distribution, corrected for multiple comparisons using Benjamini-Hochberg procedure with a false discovery rate (FDR) of *p* < 0.01. Importantly, this approach allowed the use of non-negative response profiles while permitting negative voxel coefficients, identifying cortical areas where a dimension’s profile was negatively correlated with voxel tuning. This preserved the part-based, sparse, and interpretable nature of the dimensions while relaxing the non-negativity constraint for voxel patterns. A strong linear relationship between the spatial mappings from BNMF and the encoding model confirmed that relaxing the non-negativity constraint did not compromise the validity of the original weights (mean *r* across participants ± *SD*; FFA: *r* = 0.94 ± 0.04; EBA: *r* = 0.89 ± 0.05; PPA: *r* = 0.89 ± 0.06). Noise ceilings were computed for the test set following the methodology provided in the original NSD paper (Allen et al., 2022).

### Representational sparseness

We quantified the extent of multidimensional tuning using a measure of representational sparseness based on the normalized relationship between the *L_1_*-and *L_2_*-norm of a vector (Hoyer, 2004; Eq. 5). For each voxel, the dimension weights of the encoding model were interpreted as its *n* dimensional tuning profile *x*.

The resulting sparseness index *s* ranges from 0, indicating a perfectly sparse representation tuned to a single dimension, to 1, indicating a perfectly dense representation tuned equally to all dimensions.

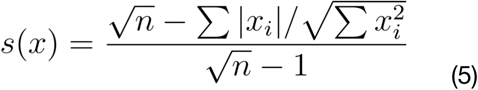

## Acknowledgments

L.E.v.D was supported by a doctoral scholarship awarded by the German Academic Scholarship Foundation. M.N.H. and K.D. were supported by the ERC Starting Grants COREDIM (ERC-2021-STG-101039712) and DEEPFUNC (ERC-2023-STG-101117441), respectively, as well as the Hessian Ministry of Higher Education, Research, Science and the Arts (LOEWE Start Professorships and Excellence Program “The Adaptive Mind”) and the Deutsche Forschungsgemeinschaft (DFG, German Research Foundation, 222641018-SFB/TRR 135 TP). The funding organizations had no role in the study design, data collection and analysis, decision to publish, or preparation of the manuscript. Computational resources were provided by the high-performance computing clusters at the Max Planck Computing & Data Facility (MPCDF), Garching, Germany.

## Author contributions

L.E.v.D., M.N.H., and K.D. conceived the study. L.E.v.D. carried out the data analysis and wrote the original draft of the manuscript. M.N.H. and K.D. reviewed the manuscript and provided critical feedback. M.N.H. and K.D. jointly supervised the project.

## Data and code availability

The data supporting our analyses were obtained from the publicly available Natural Scenes Dataset (http://naturalscenesdataset.org/). The Python code (version 3.8.20) used for data analysis and visualization is publicly available on GitHub (https://github.com/levandyck/roidims).

## Supplementary material

### Supplementary figures

**Fig. S1.**
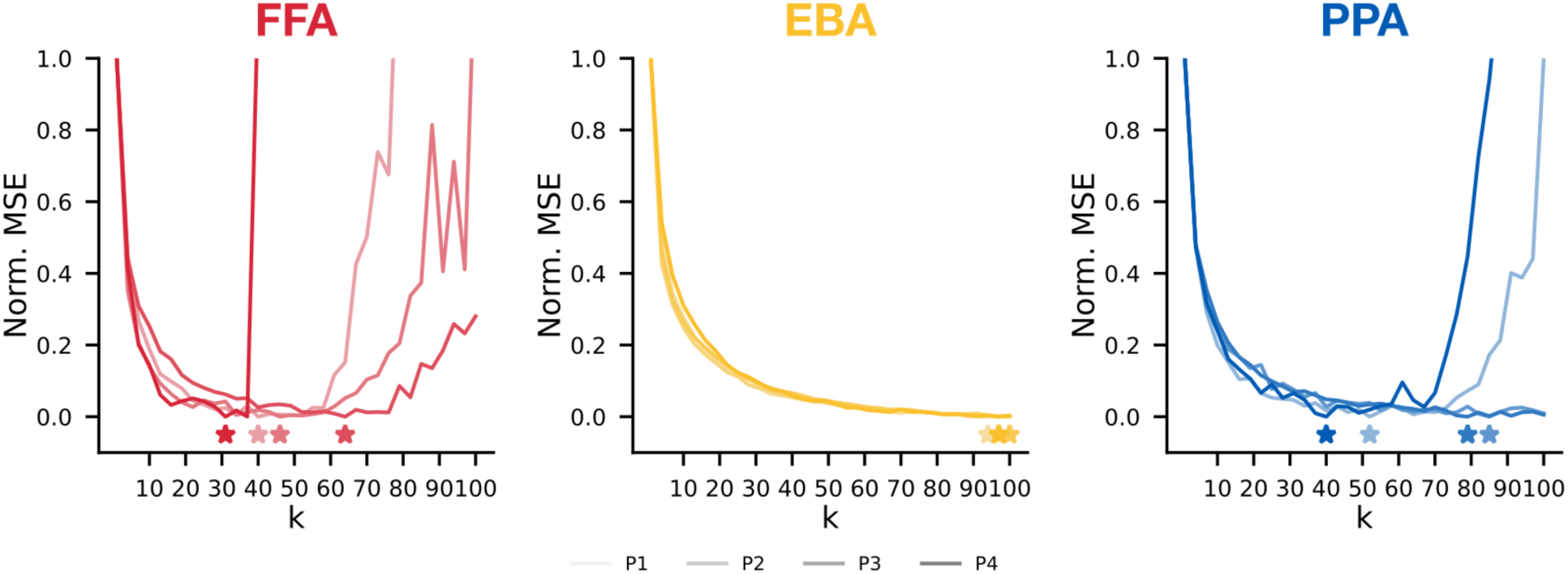
Optimal dimensionality for each area. Normalized mean squared error (*MSE*) based on bi-cross-validation for each area in each participant. Stars indicate optimal dimensionality for each participant.

**Fig. S2.**
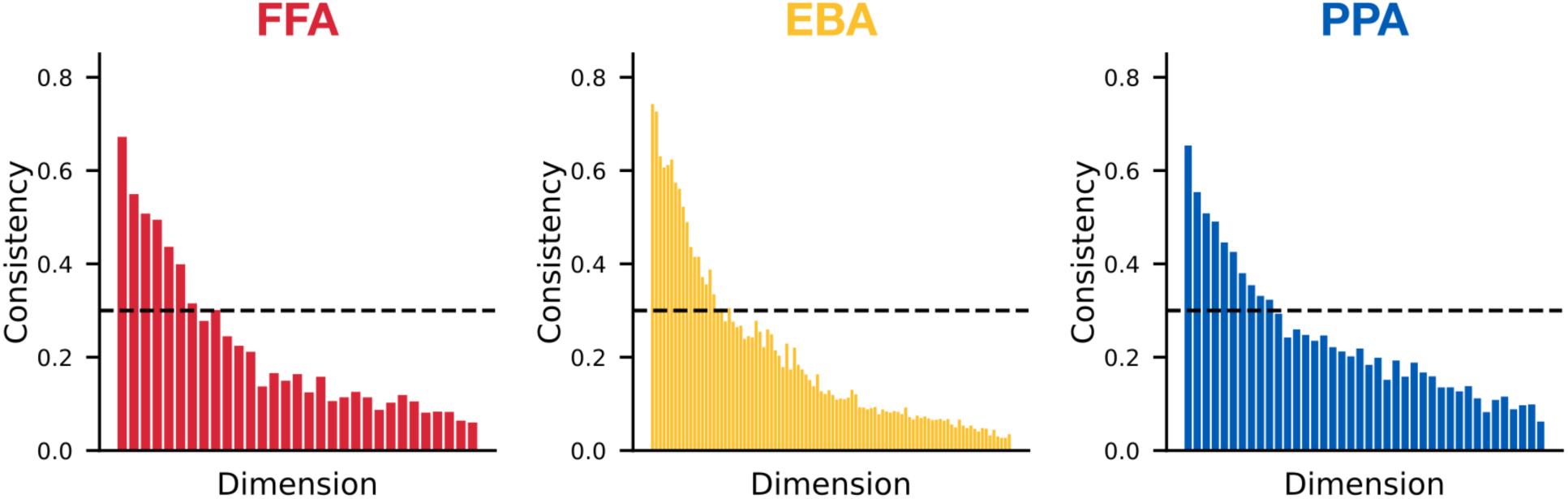
Consistency of dimensions in each area across participants. Mean correlation of matched dimensions across participant pairs. Dotted line indicates consistency threshold (*r* = 0.3).

**Fig. S3.**
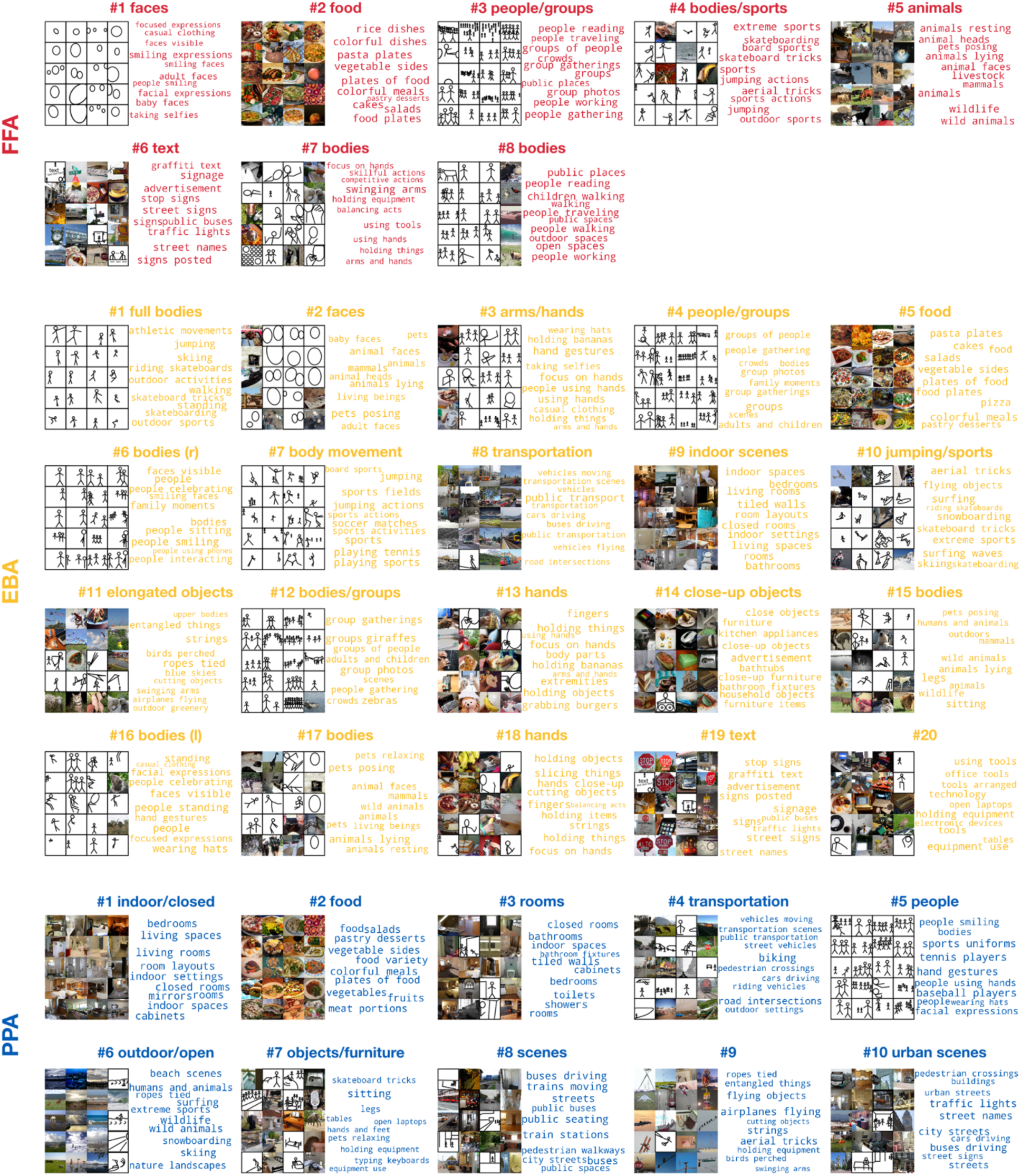
Highest-scoring images and word-clouds for individual dimensions. Five highest-scoring images per participant and ten highest-scoring labels combined across participants for individual dimensions from each area. Images of people were replaced with stick figures.

**Fig. S4.**
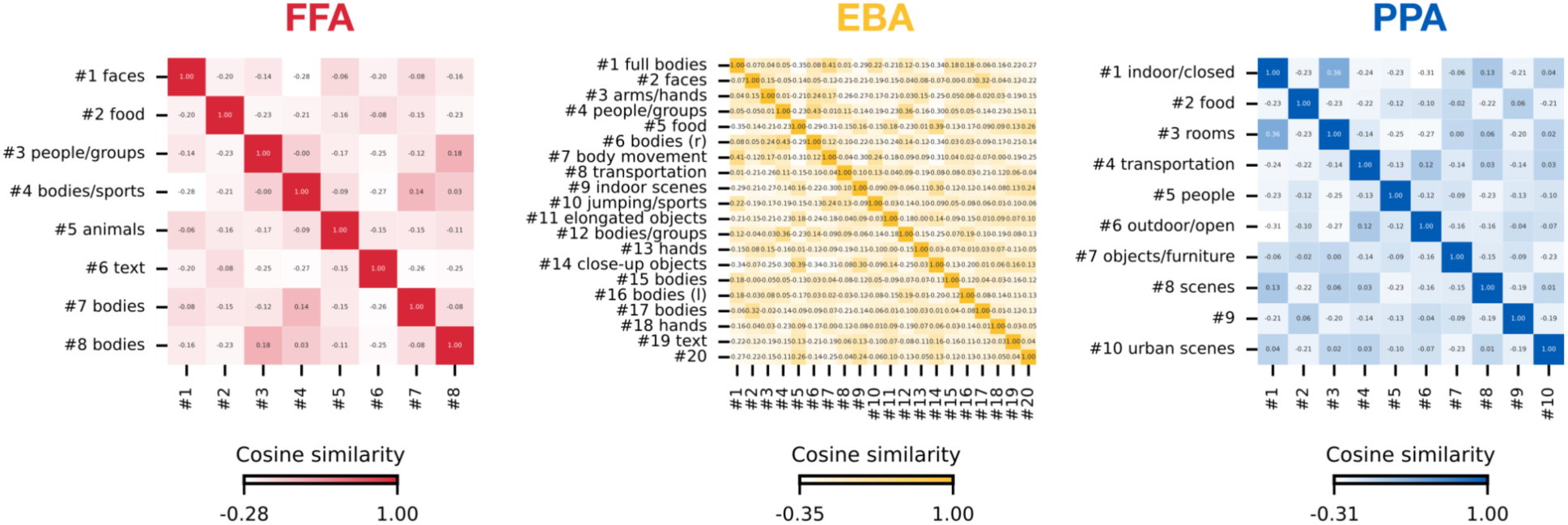
Similarity of dimensions in each area. Mean-centered pairwise cosine similarities of dimensions in each area, averaged across participants.

**Fig. S5.**
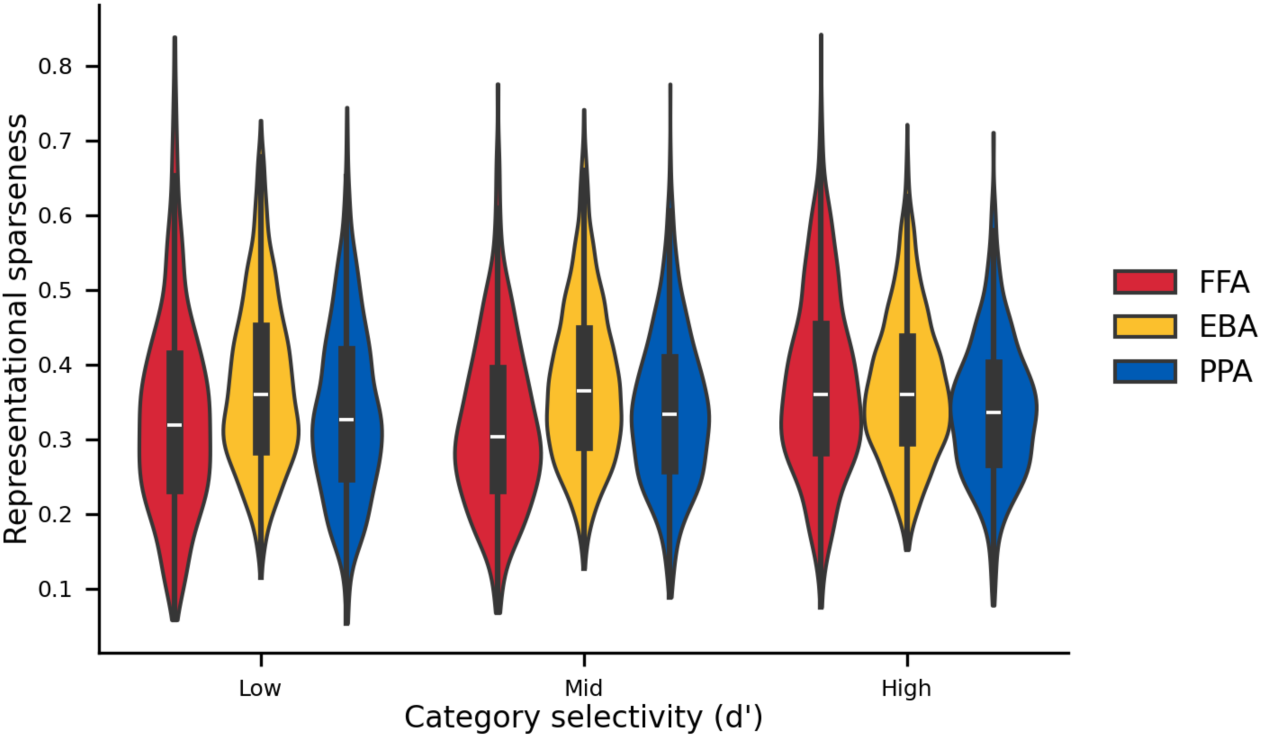
Voxel-wise representational sparseness across different levels of category selectivity in each area. Voxels within each area were grouped into low, mid, and high category selectivity (*d’*), defined using the corresponding functional localizer contrast (e.g., faces > all other categories in FFA). Representational sparseness was then computed for each voxel, with distributions shown across selectivity bins and areas.

**Fig. S6.**
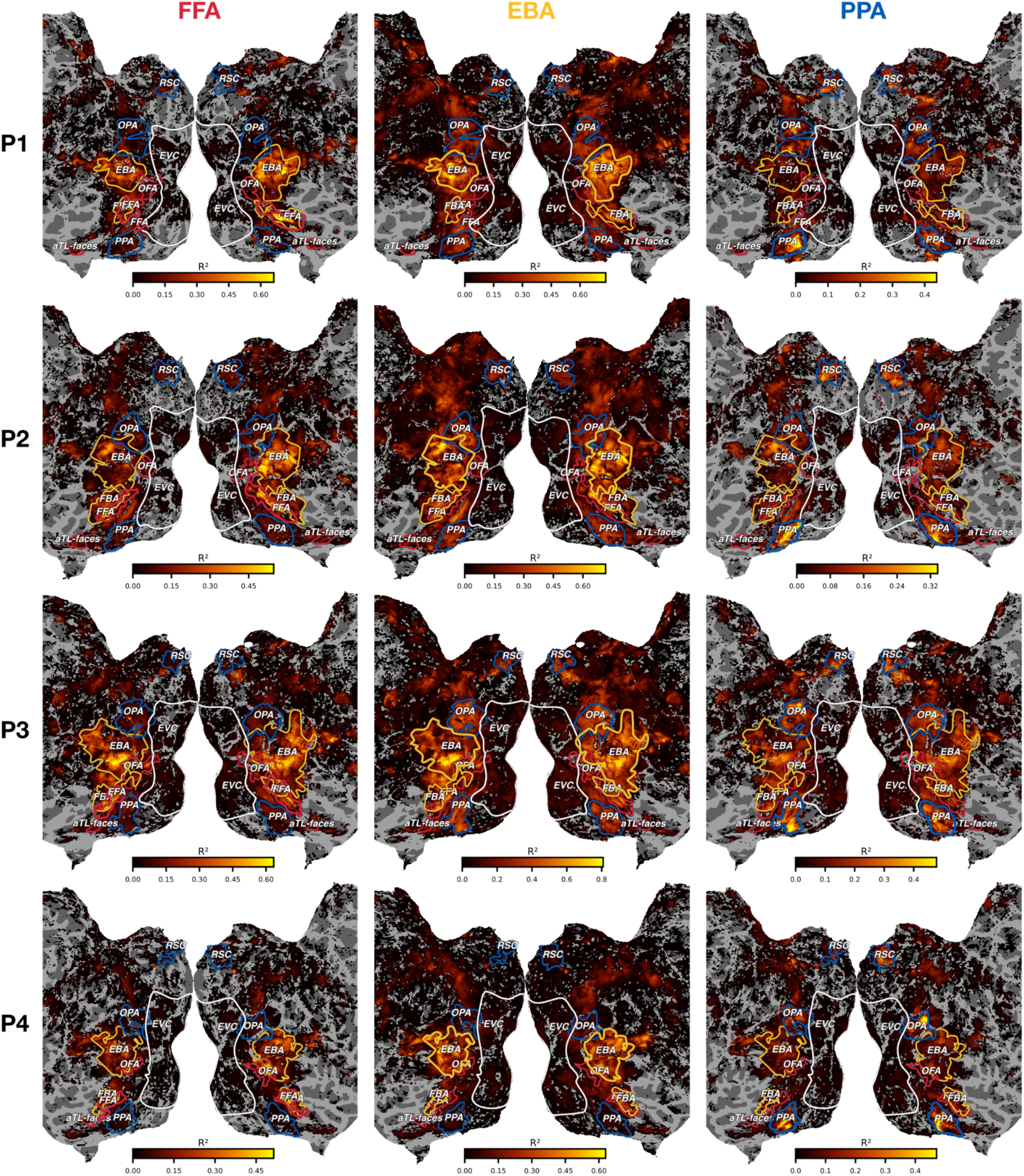
Prediction performance maps of dimensions from category-selective areas across cortex. Voxel-wise prediction performance (*R^2^*) for dimensions from each area projected onto the flattened cortical surface for all participants. Voxels thresholded at *p* < 0.01 (one-sided, FDR-corrected).

**Fig. S7.**
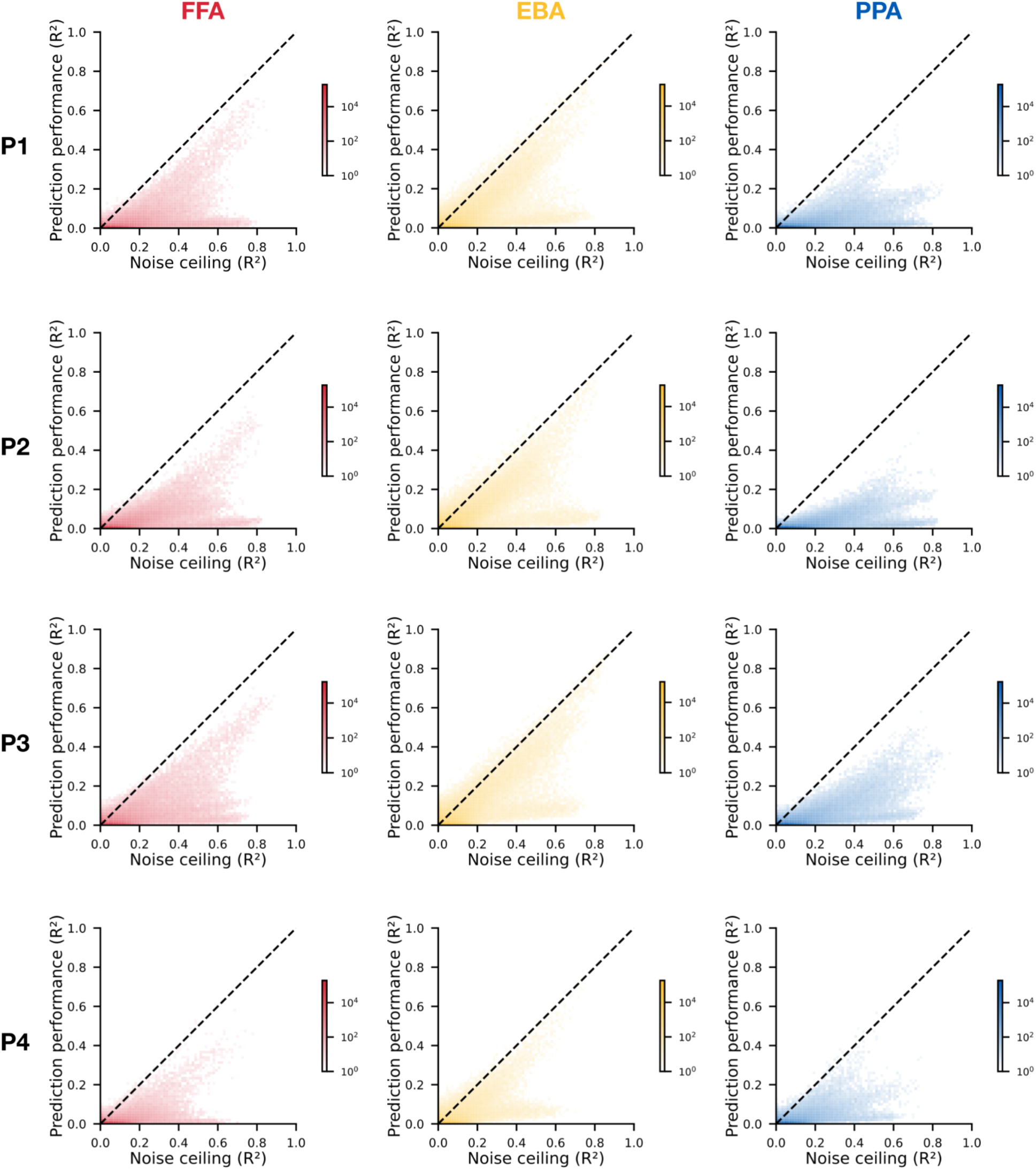
Noise ceiling estimate versus voxel-wise prediction performance across cortex. Relationship between noise ceiling estimate and prediction performance (*R^2^*) for dimensions from each area, evaluated on held-out test images. Note that both metrics exhibit variability, particularly in voxels with lower signal-to-noise ratio, likely due to the limited size of the test set (30 % of all images).

**Fig. S8.**
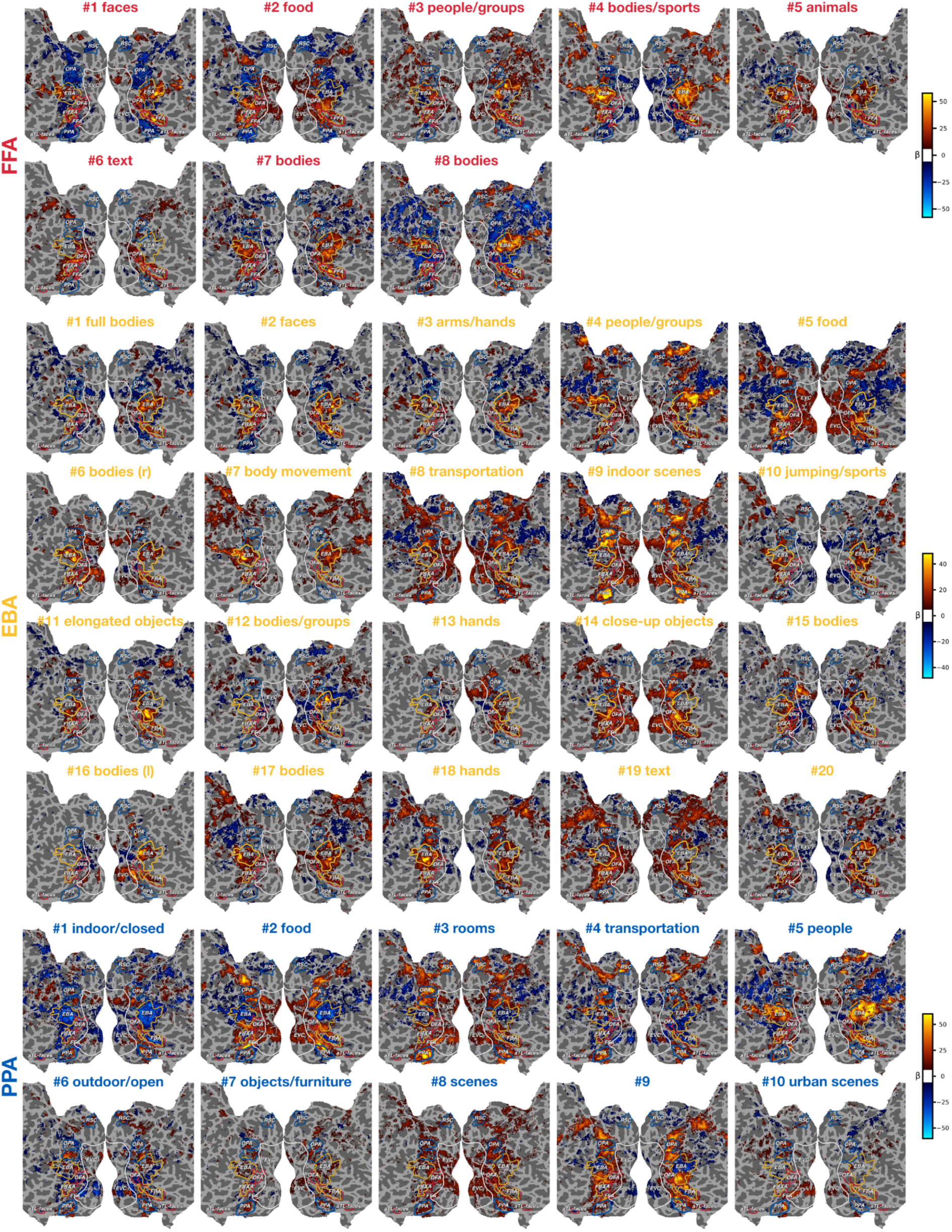
Dimension tuning maps for participant P1. Voxel-wise coefficient weights for individual dimensions from each area projected onto the flattened cortical surface. Voxels thresholded at *p* < 0.01 (one-sided, FDR-corrected).

**Fig. S9.**
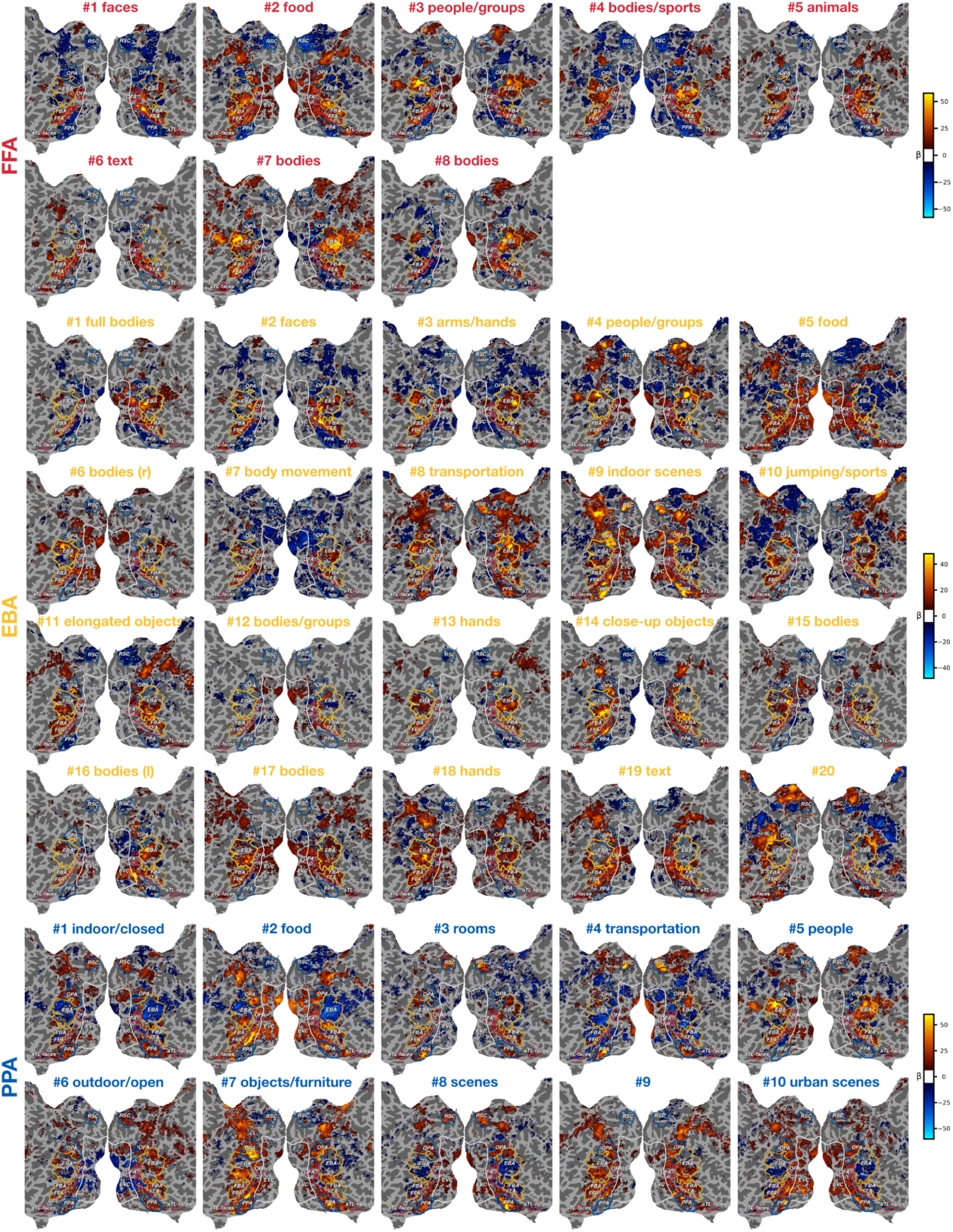
Dimension tuning maps for participant P2. Voxel-wise coefficient weights for individual dimensions from each area projected onto the flattened cortical surface. Voxels thresholded at *p* < 0.01 (one-sided, FDR-corrected).

**Fig. S10.**
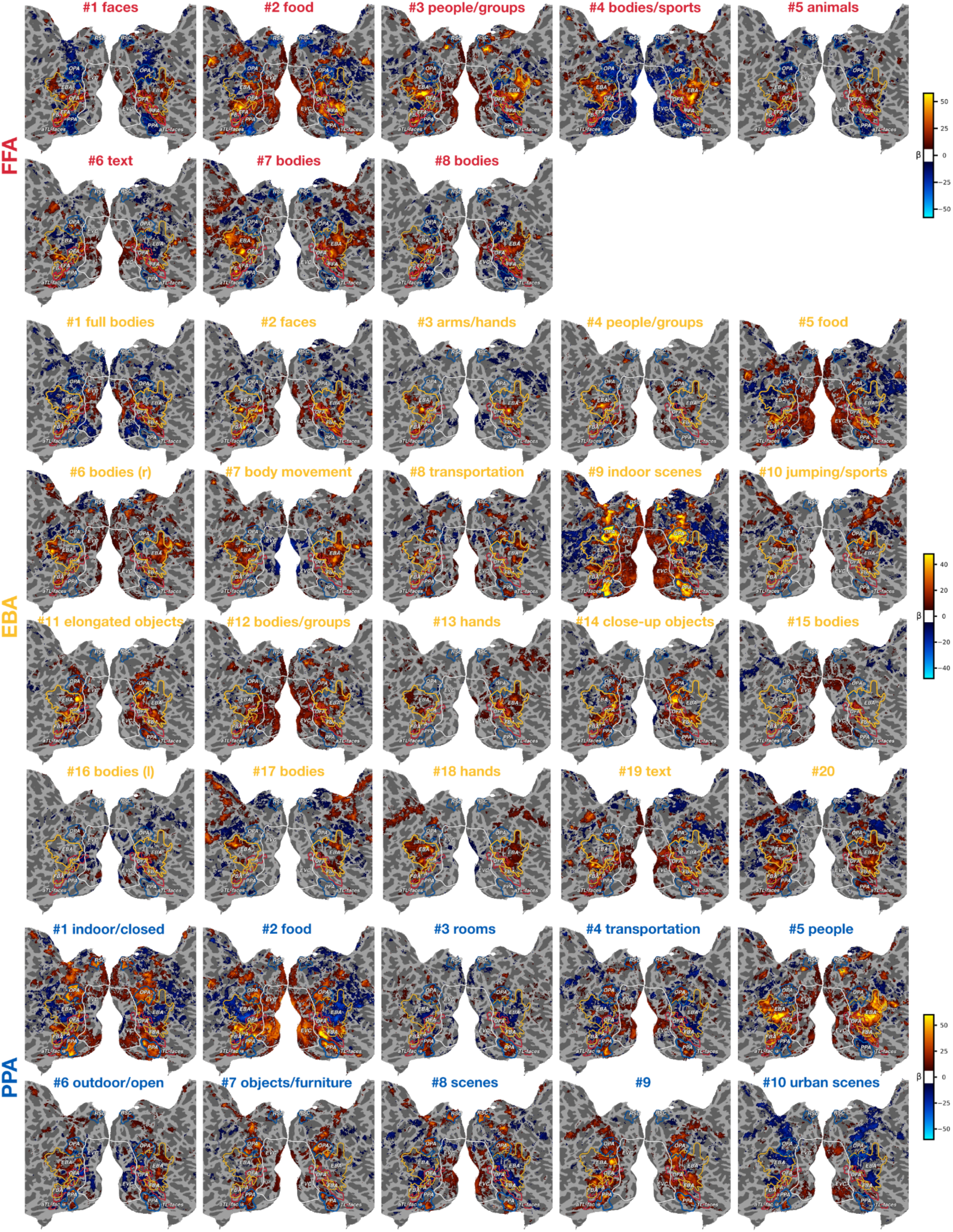
Dimension tuning maps for participant P3. Voxel-wise coefficient weights for individual dimensions from each area projected onto the flattened cortical surface. Voxels thresholded at *p* < 0.01 (one-sided, FDR-corrected).

**Fig. S11.**
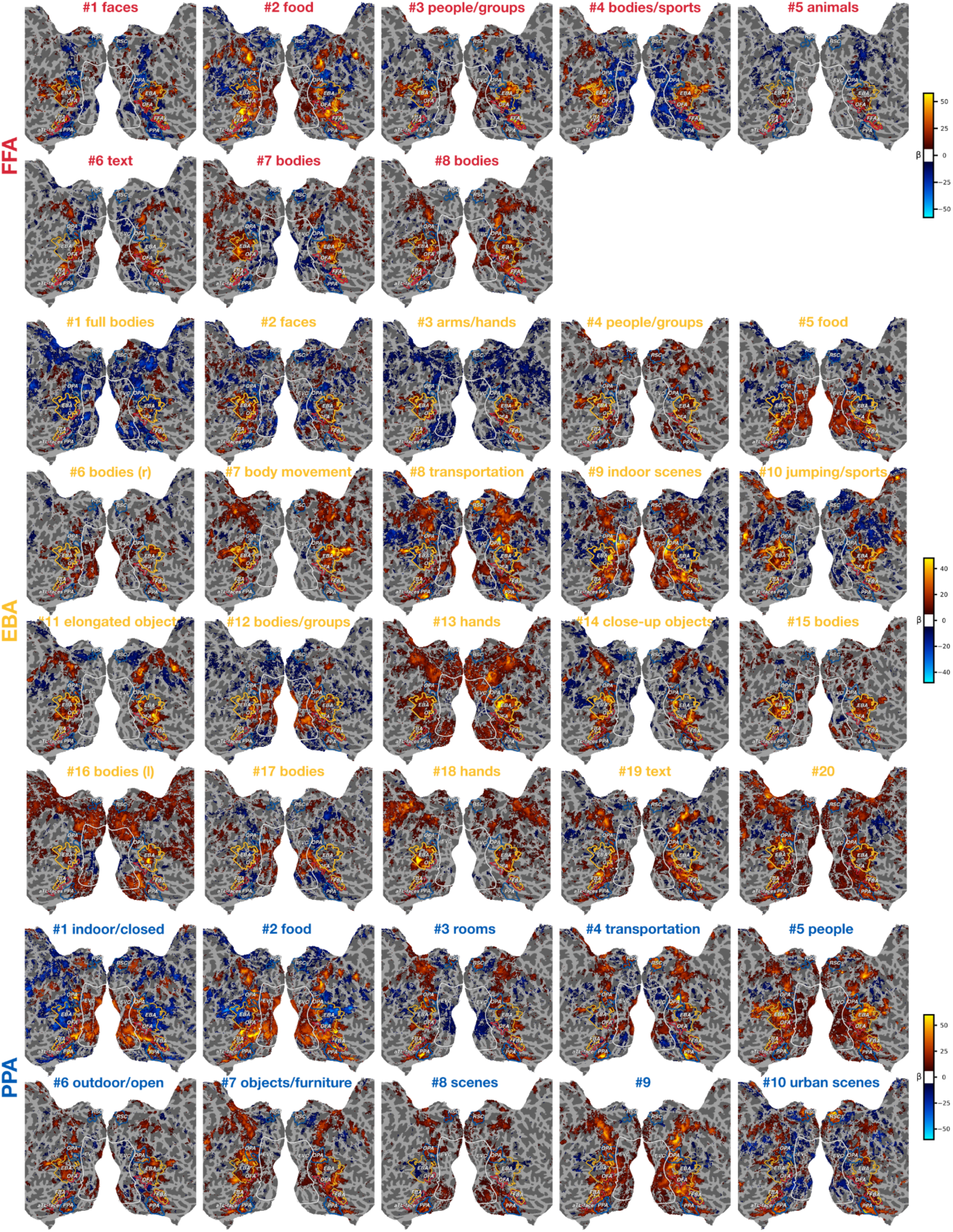
Dimension tuning maps for participant P4. Voxel-wise coefficient weights for individual dimensions from each area projected onto the flattened cortical surface. Voxels thresholded at *p* < 0.01 (one-sided, FDR-corrected).

**Fig. S12.**
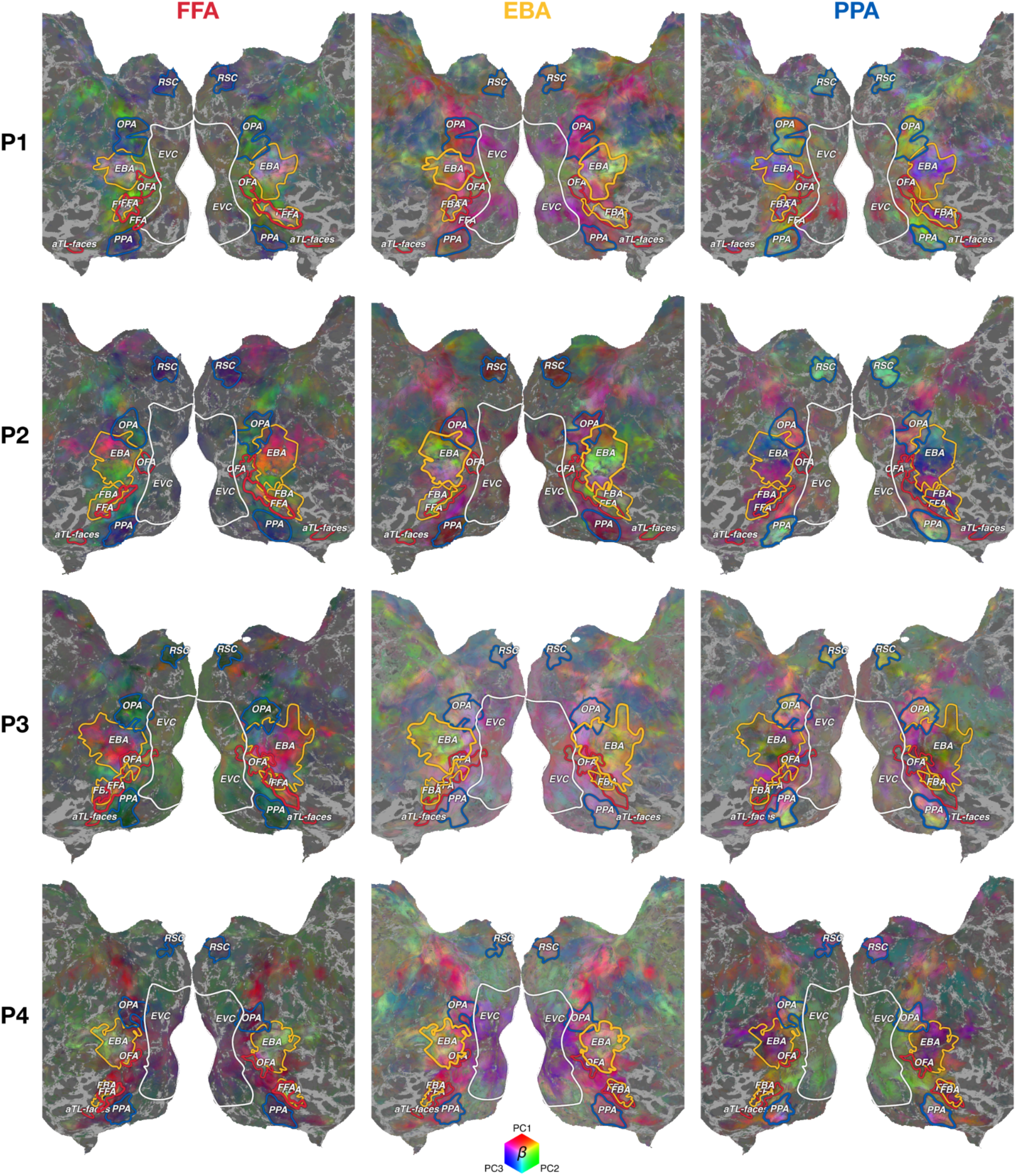
Functional tuning gradients based on all dimensions. First three principal components (PC1-3; red, green, and blue channels) of voxel-wise coefficient weights across all dimensions projected onto the flattened cortical surface for all participants. Voxels thresholded at *p* < 0.01 (one-sided, FDR-corrected).

### Supplementary text

We used GPT-4V with the following prompt to generate candidate labels based on the highest-scoring images: *Analyze the following collage of images and identify 10 short labels that describe the most prominent, recurring elements observable across all or nearly all images*.

*Each label should:*

-*Focus solely on common visual features (e.g., objects, categories, scenes)*.
-*Consist of up to three specific and concrete words (preferably less; preferably nouns and verbs)*.
-*Match the sentence:’The images show {label}.*’

